# An Integrated Experimental and Informatics Strategy Enables Large Proteoform Identification in Top-Down Proteomics

**DOI:** 10.64898/2026.10.30.763218

**Authors:** Matthew S. Fischer, Eli J. Larson, Ryan Schroeder, Hsin-Ju Chan, Heewon Choi, Thomas S. Weir, Andrew J. Perciaccante, Carmen C. Sucharov, Yang Lu, Ying Ge

## Abstract

Top-down proteomics enables identification and characterization of intact proteoforms, preserving information on co-occurring sequence variations and post-translational modifications. However, identifying large proteoforms remains challenging due to an exponential decrease in signal-to-noise ratio with increasing molecular weight, interference from coeluting smaller species, unresolved precursor isotopic envelopes, difficulty obtaining informative MS/MS spectra, and limitations of current informatics tools. To address these challenges, here we present an integrated experimental and informatics strategy combining protein separation, data acquisition, precursor mass assignment, and MS/MS interpretation. First, we demonstrate that online two-dimensional liquid chromatography coupling size-exclusion and reversed-phase separations enables facile detection of large proteoforms. We then utilize a precursor mass correction strategy to enable conventional database searching of data-dependent acquisition (DDA) data containing isotopically unresolved precursors. Finally, we use data-independent acquisition (DIA) optimized for large proteins to enhance fragment signal and enable identification through intact-mass-guided candidate discovery and MS/MS validation. Applied to human heart tissue, the integrated strategy identified 703 nonredundant proteoforms from 236 unique proteins, including 65 proteins >30 kDa and 22 proteins >50 kDa, with identifications extending to 223 kDa. This integrated strategy bridges the detection-to-identification gap for large proteins and expands the accessible mass range of discovery top-down proteomics.

## Introduction

Proteoforms are distinct molecular forms of proteins arising from combinations of genetic variation, alternative splicing, and post-translational modifications that contribute to the structural and functional diversity of proteins in biological processes and diseases.^1–3^ Mass spectrometry-based top-down proteomics analyzes intact proteins without enzymatic digestion, enabling proteoform identification and characterization while preserving information on co-occurring sequence variations and modifications.^4–9^ In recent years, mass spectrometry-based top-down proteomics has benefited from rapid advancement in mass spectrometry (MS) instrumentation, intact protein separations, tandem mass spectrometry methods, and informatics strategies which have increased the number of confident proteoform identifications achievable from biological samples.^10–14^

Despite technological advancement, the identification of larger proteoforms (>30 kDa) in discovery top-down proteomics workflows remains challenging.^6,15^ As the molecular weight of a protein increases, the increasing number of charge states and isotopologues results in an exponential decrease in signal-to-noise ratio.^16^ Furthermore, the increasing number of bond cleavage sites decreases fragment ion intensity and increases spectral complexity. Consequentially, discovery top-down studies are dominated by small proteoforms.^17–21^ For example, in one of the largest top-down studies to date, nearly 90% of the 20,000+ identified proteoforms were under 10 kDa,^19^ and in some protocols larger proteins (>30 kDa) are excluded during sample preparation altogether.^22–25^ Given that more than 70% of proteins in the UniProt human reference proteome have molecular weights greater than 30 kDa (∼15% are >100 kDa),^26^ strategies to extend the mass range of discovery top-down proteomics are urgently needed.

Protein separation is a critical first step toward addressing this coverage gap. Size-based prefractionation methods can facilitate detection of large proteins by minimizing coelution with smaller proteins, but a detection-to-identification gap persists. For example, a previous method using offline serial size-exclusion chromatography (SEC) followed by reversed-phase liquid chromatography (RPLC) enabled the detection of large proteoforms (up to 200+ kDa) yet only reported MS/MS identification of two proteins >30 kDa.^27^ Similarly, despite the use of gel-eluted liquid fraction entrapment electrophoresis (GELFrEE) fractionation, 93% of proteoforms identified for the Blood Proteoform Atlas are smaller than 20 kDa.^17^

Informatics limitations present an additional barrier.^4^ Identification of large proteins from complex mixtures often requires numerous targeted experiments^28–30^ and multiple identification methods which are difficult to incorporate into automated discovery pipelines. Conventional search tools generate proteoform-spectrum matches by comparing individual LC-MS/MS spectra with database sequence ladders.^31,32^ Many top-down proteomics software tools including TopPIC Suite,^33^ pTOP,^34^ and Informed-Proteomics^35^ rely on isotopically-resolved data for both deconvolution and searching, but isotopic resolution is difficult to achieve for large proteins. As a result, incorrect precursor masses are assigned to isotopically unresolved proteins, precluding identification. Alternatively, Schaffer *et al.* demonstrated Proteoform Suite for the identification of large proteins by intact average mass, but this strategy suffered from a high false discovery rate (FDR) and was ultimately used for generating plausible candidates for targeted MS/MS validation, resulting in identification of only a small fraction of the large proteoforms detected.^36^

To address these limitations, we have designed and developed an integrated experimental and informatics strategy for large protein identification in discovery top-down proteomics. First, we develop an online 2D-LC-MS/MS method coupling microscale SEC and RPLC (**Figure 1A, B**). This online 2D-LC approach provides a higher peak capacity than a conventional one-dimensional RPLC separation while reducing the labor and sample requirements of offline workflows, enabling the detection of numerous large proteoforms from human heart tissue. To address the large protein detection-to-identification gap, we developed two complementary identification strategies. The first method is an algorithm for precursor mass correction, allowing large proteins lacking isotopic resolution to be identified from data-dependent acquisition (DDA) by a conventional database searching tool, TopPIC Suite (**Figure 1C**). The addition of a precursor mass correction step increased the number of unique identified proteins >30 kDa from 3 to 41. However, for large proteoforms, signal distribution across multiple precursor charge states limited fragment ion signal in individual DDA spectra. Therefore, we developed a second method involving data-independent acquisition (DIA) tailored to large protein fragmentation with intact-mass-guided candidate discovery and MS/MS validation (**Figure 1D**), identifying an additional 24 proteins averaging approximately 73 kDa. Overall, this integrated workflow identified 703 proteoforms from 236 unique proteins from human heart tissue, including 65 unique proteins with masses greater than 30 kDa. This corresponded to 28% of identified proteins and 18% of identified proteoforms having masses over 30 kDa, extending to a maximum mass of 223 kDa. These advances significantly expand high molecular weight protein coverage in discovery top-down proteomics.

**Figure 1.**
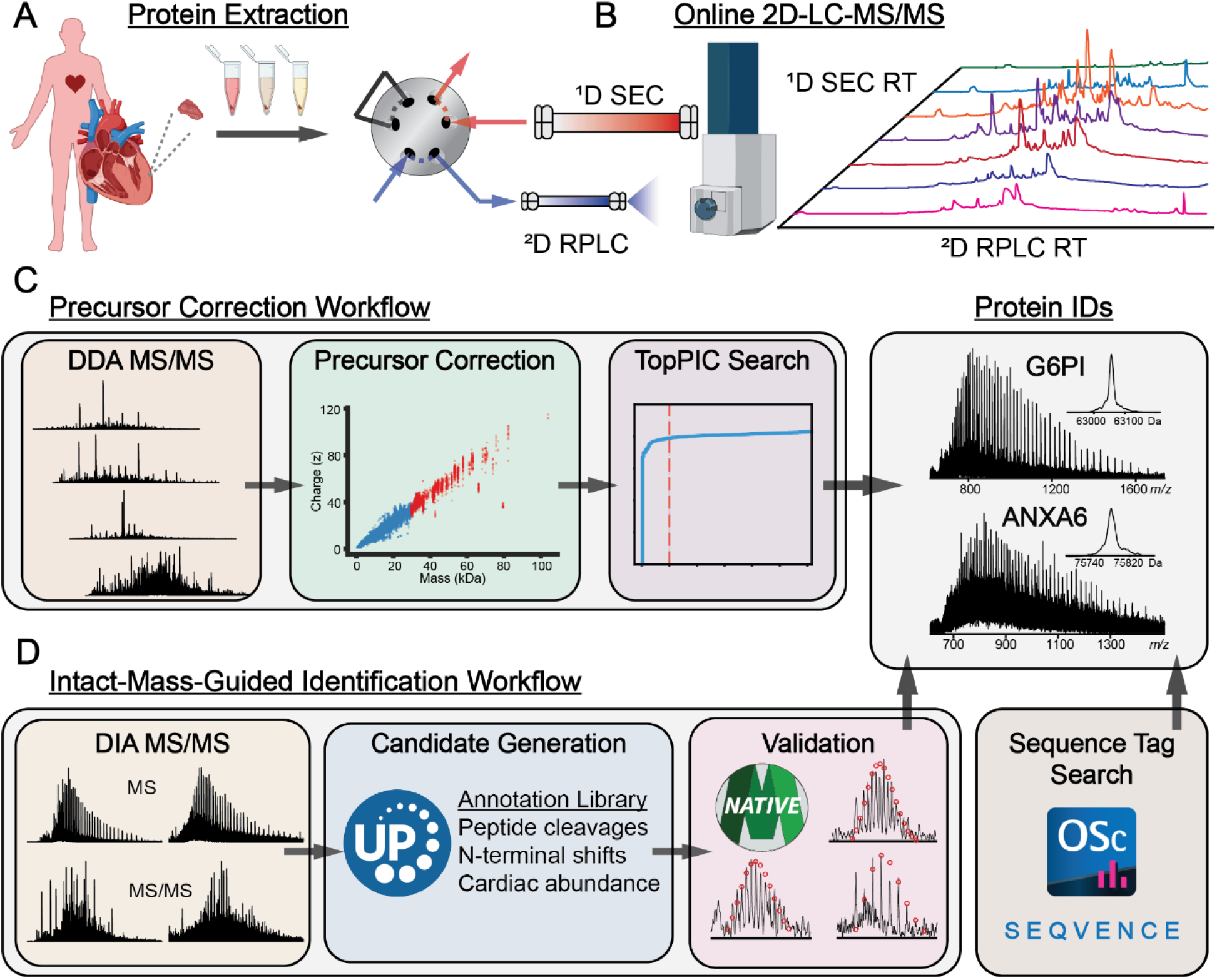
Integrated experimental and informatics framework for large protein identification. **A)** Sequential extraction of human heart tissue using HEPES buffer, trifluoroacetic acid (TFA), and the photocleavable surfactant Azo generates fractions enriched in soluble, contractile, and membrane-associated/insoluble proteins, respectively. **B)** Online two-dimensional liquid chromatography-tandem mass spectrometry (2D-LC-MS/MS) combines size-based fractionation with reversed-phase separation to reduce sample complexity and facilitate detection of large proteins. **C)** For data-dependent acquisition (DDA), UniDec-based precursor mass correction replaces incorrect mass assignments for isotopically unresolved precursors. Corrected precursor masses are paired with the corresponding MS/MS fragment lists for TopPIC database searching. **D)** Data-independent acquisition (DIA) samples multiple precursor charge states to enhance fragment signal. Intact-mass-guided candidate discovery compares measured average masses with an annotated human-heart protein library, considering sequence processing and common N-terminal modifications. Candidates are ranked by mass agreement and protein abundance, followed by MS/MS validation in MASH Native. A separate complementary route uses Bruker OmniScape to generate *de novo* sequence tags for MS-BLAST searching when intact-mass-guided candidate discovery does not yield a verifiable identity.

## Experimental Section

### Chemicals and Reagents

All reagents were purchased from Millipore Sigma (Burlington, MA, USA) unless otherwise noted. Solutions were prepared with LC-MS-grade water from Milli-Q (Millipore, Corp., Billerica, MA, USA). Isopropanol (IPA), acetonitrile (ACN) and hexafluoroisopropanol (HFIP) were purchased from Fisher Scientific (Fair Lawn, NJ, USA).

### Human Cardiac Tissue Collection

Healthy donor hearts with no history of heart disease were obtained from the University of Wisconsin–Madison Organ and Tissue Donation-Surgical Recovery and Preservation Services. Tissue was stored in cardioplegic solution until dissection and then flash-frozen in liquid nitrogen and stored at −80 °C. The procedures for the collection of human donor heart tissues were approved by the UW–Madison Institutional Review Board.

### Protein Extraction from Human Heart Tissue

Protein was extracted following a 3-step sequential extraction method developed by Gao et al.^37^ Approximately 25 mg of human heart tissue was homogenized in 150 μL of HEPES extraction buffer (25 mM HEPES (pH = 7.4), 60 mM NaF, 2 mM EDTA, 1 mM L-Met, 1 mM TCEP, 1 mM PMSF, 1x HALT protease/phosphatase inhibitor cocktail) using a Teflon pestle in a 1.5 mL low-bind microcentrifuge tube at 4 °C. The homogenate was centrifuged for 30 min at 21,000 g at 4 °C. The supernatant was removed and saved as the soluble “HEPES” protein fraction. The remaining pellet was homogenized in 150 μL of TFA extraction solution (1% TFA, 5 mM L-Met, 1 mM TCEP) and centrifuged for 30 min at 21,000 g at 4 °C. The supernatant was removed and saved as the sarcomeric “TFA” fraction. Finally, the remaining pellet was homogenized in 150 μL of Azo extraction buffer (0.4% 4-hexylphenylazosulfonate (Azo), 25 mM ammonium bicarbonate pH 8.0, 10 mM L-Met, 5 mM TCEP, 5 mM EDTA, 1 mM PMSF, and 1x HALT Protease and Phosphatase Inhibitor Cocktail) and centrifuged for 30 min at 21,000 g at 4 °C. The supernatant was removed and saved as the insoluble “Azo” fraction. The HEPES and TFA fractions were exchanged into the ^1^D mobile phase composition using an Amicon 10 kDa molecular weight cut-off filter. The Azo fraction was subjected to a chloroform-methanol-water precipitation and resolubilization according to a previously described method.^38^ A Bradford protein assay was used to determine the final concentration of all fractions prior to analysis.

### Online 2D-LC-MS/MS Analysis of Human Heart Extracts

All liquid chromatography–mass spectrometry experiments were conducted with a Waters ACQUITY UPLC M-Class and Bruker maXis II quadrupole time-of-flight mass spectrometer. For SEC, the separation column was a PolyHYDROXYETHYL A™ column (2.1×200 mm, 5 μm, 1000 Å) from PolyLC Inc. (Columbia, MD, USA). The RPLC column was a home-packed capillary (0.5×100 mm) containing 2.7 μm, 1000 Å HALO C4 material purchased from Advanced Materials Technologies (Wilmington, DE, USA).

The ^1^D mobile phase contained 1% formic acid, 0.1% hexafluoroisopropanol, and 10% IPA in water, with a flow rate of 30 μL/min. The ^2^D mobile phases were 0.2% formic acid in water (A) and 0.2% formic acid in ACN (B). The ^2^D flow rate was 24 μL/min, and the column was heated to 60 °C. The valve interface was set up such that a valve switch allowed ^1^D effluent to flow directly into the ^2^D column, trapping eluting proteins for the duration of fraction collection, which were then separated with a gradient following reversal of the valve switch (**Figure S1**). Up to seven 1-min fractions were transferred from the first dimension depending on the extract analyzed, with SEC elution times between 11 and 18 minutes. The ^2^D gradient was set to begin at the end of fraction collection and followed this program: 10-65% B in 60 min, 65-100% B in 5 min. Before analysis of a fraction, the ^2^D column re-equilibrated at 10% B. For HEPES and TFA extracts, 25 μg of protein was injected, and 8 μg of protein was injected from Azo extracts.

For MS analysis, the source and tune parameters were as follows: end plate offset of 500 V, capillary voltage of 4500 V, nebulizer pressure of 1 bar, dry gas flow rate of 5 L/min, dry gas temperature of 220 °C, funnel 1 RF of 400 Vpp, isCID energy of 10 eV, multipole rf of 300 Vpp, quadrupole ion energy of 10 eV, low mass cutoff at 300 *m/z*, collision cell energy of 4 eV, collision RF of 2500 Vpp, transfer time of 115 μs, and pre pulse storage of 20 μs. For DDA MS/MS, the top 3 precursors were selected for fragmentation with active exclusion enabled for 0.5 min after 2 spectra, with collision energy according to **Table S1**. The scan rate for MS and MS/MS was 1 Hz. For DIA MS/MS, fragmentation was performed in two windows, 800-900 *m/z* (12 eV) and 900-1000 *m/z* (18 eV). Acquisition was in the order of a single MS scan (1 Hz) followed by a scan for each MS/MS windows (0.5 Hz). For the identification of Myosin-7, a different method was used which isolated a window ranging from 1000-1100 *m/z* (15 eV).

### Precursor Mass Correction and Database Searching

Raw Bruker data files were centroided and converted to *.mzML using MSConvert (Proteowizard 3.0).^39^ The centroided *.mzML files were deconvoluted using TopFD,^40^ producing output in the *.msalign format. Because TopFD is designed for only isotopically resolved data, we implemented UniDec^41^ 8.1 in Python to correct the masses of unresolved precursors. Our algorithm implemented the following procedure: 1) each MS1 scan is deconvoluted with 1 Da resolution, 2) precursor masses are calculated over a range of reasonable charge states (5 to 120) and matched to UniDec deconvolution results within an intensity threshold (10% of scan maximum) and mass error window (100 ppm), 3) a new *.msalign file is generated consisting only of scans where precursor masses were replaced by UniDec values. For a matched mass to be written, it was required to have been isolated at least twice, been present in a scan with no greater than 15 UniDec picked peaks and exceed a mass threshold (30 kDa). This pipeline was only applied to the early-eluting SEC fractions which contained large proteins. The *.msalign outputs from TopFD and the precursor mass correction algorithm were searched with TopPIC 1.8 at 1% protein and proteoform level FDR against the UniProt reviewed human proteome FASTA (canonical and isoform).^33^ Precursor mass values for one data file were also compared against results from FLASHDeconv^42^ (OpenMS 3.5) on profile data.

### Intact-Mass-Guided Candidate Discovery and MS/MS Validation

MS/MS spectra corresponding to unidentified large proteins were summed from DIA results. A candidate list was assembled for each unidentified protein average mass using a database consisting of human heart proteins identified previously by bottom-up proteomics.^43^ Detected masses were matched against the database proteins according to their UniProt annotated form (including transit/signal peptide cleavages and N-terminal modifications). Mass shifts corresponding to common N-terminal alterations (**Table S2**) were also considered as variable shifts. Candidates were sorted into three groups based on their mass shifts: 1) “exact matches” with theoretical masses within 2 Da of the measured mass; 2) “shift rescues”, for which a common N-terminal variable shift that is not annotated on UniProt brings the mass within 2 Da, and 3) “vicinity candidates”, which was set to be the nearest masses within 500 Da, with a maximum of 10 candidates. Within each group, candidates are sorted by their abundance according to previous bottom-up proteomics results.^43^ Plausible candidates were validated against their respective MS/MS spectra in MASH Native.^44^ Candidates were accepted if their sequences resulted in a sufficient number of matching terminal fragments within 15 ppm error tolerance relative to overall spectral complexity (the number of matching fragments for accepted candidates ranged from 5 to 82, with an average of 27). For proteins which did not produce verifiable candidates, *de novo* sequence tags were generated with the Bruker Omniscape *de novo* pipeline and searched with MS-BLAST.^45^ When collapsing and comparing unique/redundant proteins across identification methods, truncated proteoforms were given lower priority than full-length proteoforms. For example, if a truncated proteoform was identified from a TopPIC search and the corresponding full-length proteoform is identified from a candidate list, the unique protein is considered identified by candidate validation to more accurately compare. The total number of unique proteoforms was determined by filtering according to the TopPIC ouput “Proteoform” column which contains all sequence and modification information and then manually adding and verifying the uniqueness of proteoforms identified by the intact-mass-guided workflow.

## Results and Discussion

### Development of an Online SEC-RPLC-MS/MS Method for Top-Down Proteomics

2D-LC can substantially increase peak capacity and reduce sample complexity for top-down proteomics, but most methods use offline prefractionation, which can be time-consuming, prone to sample loss, and limit throughput in discovery top-down proteomics.^27,46–51^ Online 2D-LC reduces manual handling and facilitates automation, with demonstrated improvements in sensitivity, sample requirements, and analysis time.^18,52–55^ Here, we developed an online 2D-LC method directly coupling size separation with reversed-phase separation to facilitate MS/MS fragmentation and identification over a wide range of protein molecular weights. Previously, we have shown that serial coupling of SEC columns with varying pore sizes (sSEC) can provide effective size separation of proteoforms by extending the molecular weight range of the fractionation step.^27,56,57^ Here, to improve sensitivity and focus primarily on large proteins, we selected a PolyHYDROXYETHYL A™ SEC column with a uniform pore size of 1000 Å. By performing online SEC-RPLC, we minimize sample loss and artifactual modifications which may occur during offline sample handling. Our online 2D-LC method is designed such that the highly aqueous ^1^D SEC effluent may flow directly into the ^2^D RPLC column while a size fraction is collected, trapping proteins until the start of the gradient (**Figure S1**).

We applied our SEC-RPLC-MS/MS method to a three-step sequential extraction from human heart tissue. This extraction produces three distinct fractions: a soluble protein fraction obtained with the HEPES extraction, a fraction with enriched contractile proteins from the TFA extraction, and an “insoluble” protein fraction solubilized by a photo-cleavable surfactant, Azo.^37,58^ No detectable protein elution from the RPLC column was observed during interdimensional sample transfer for all extraction conditions, indicating that proteins are sufficiently retained by the ^2^D column regardless of minor solvent mismatch. Using the TFA extraction condition as an example, the total ion chromatograms (TICs) display heterogeneity between size fractions, indicating effective separation in the first dimension (**Figure 2A**). Representative extracted ion chromatograms (EICs) and spectra from each SEC fraction of the TFA extraction conditions demonstrate coverage over a wide range protein molecular weight (<10 kDa to >200 kDa) (**Figure 2B, C**). Similar attributes were observed in the HEPES extraction condition (**Figure S2**). The peak capacity of the online 2D-LC separation was estimated to be approximately 424 following correction for undersampling of the first dimension.^59,60^ This value was obtained by generating a total of 24 EICs across various ^1^D fractions to estimate ^2^D peak capacity (**Table S3**), while the ^1^D SEC peak capacity was estimated using the method of Hagel.^61^ Overall, this represents a significant improvement over one-dimensional RPLC methods in terms of peak capacity and accessible molecular weight range.

**Figure 2.**
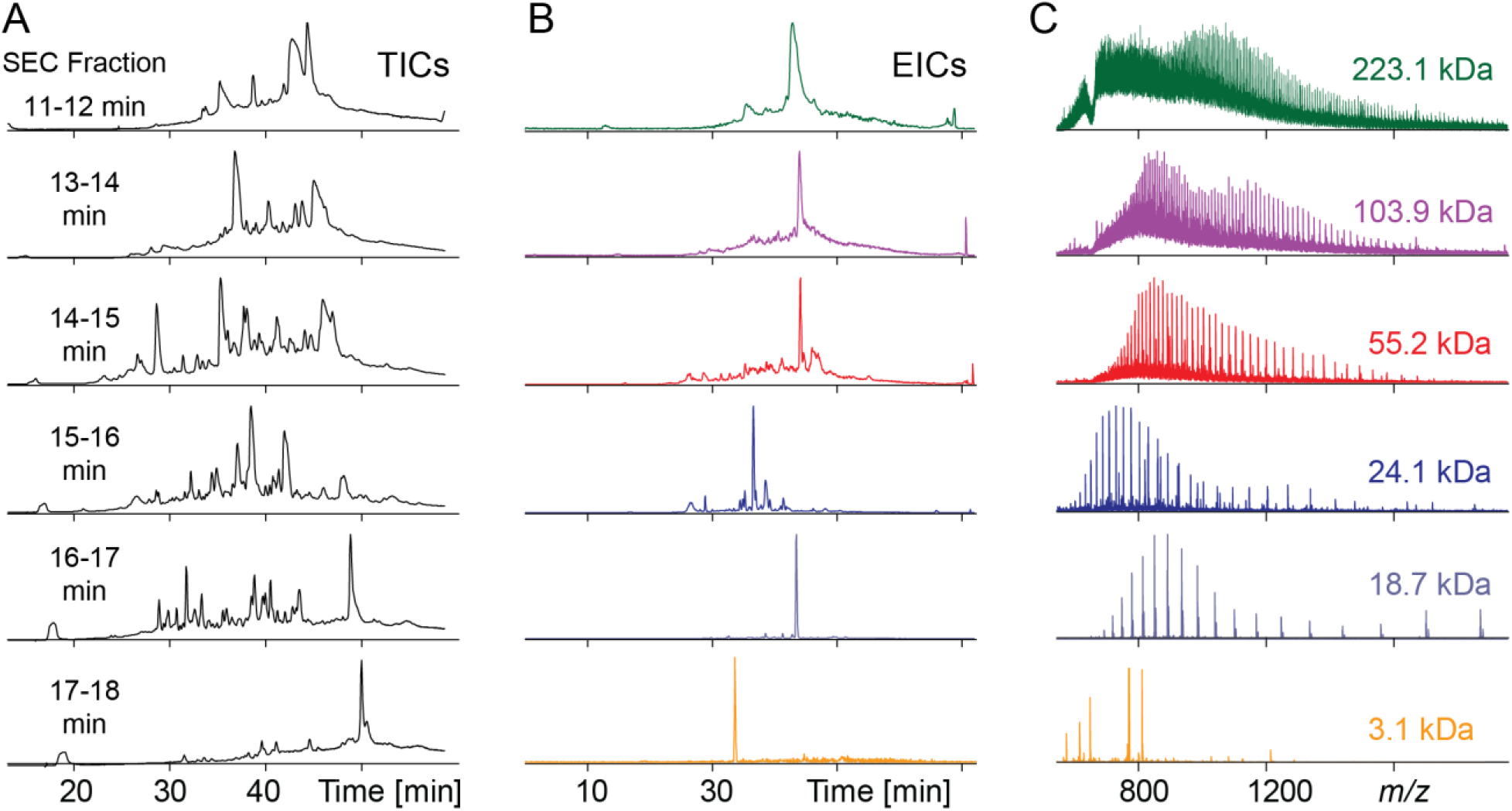
Online SEC–RPLC-MS/MS enables detection of human heart proteins across a broad molecular-weight range. **A)** Total ion chromatograms (TICs) corresponding to the RPLC separation of SEC fractions from the TFA extraction condition. The TICs show significant heterogeneity between fractions, indicating effective size separation. **B)** Representative extracted ion chromatograms and **C)** corresponding mass spectra of proteins detected within each SEC fraction, demonstrating detection over a wide range of molecular weights.

### Precursor Mass Correction Enables Identification of Large Proteins from a Database Search

To identify proteins from the 2D-LC data, we first performed a conventional database search using TopPIC Suite,^33^ deconvoluting each data file using TopFD. However, since TopFD relies on isotopically spectra, the precursor masses for large, unresolved proteins were assigned incorrectly.^40^ This prevents identification as the incorrect precursor mass will be distant from the matching sequence ladder. As a result, the initial search only identified 3 unique proteins with masses greater than 30 kDa. To enable identification of larger proteins, we developed an algorithm which performs a separate deconvolution of each MS scan with UniDec and matches the results against corresponding precursor *m/z* values at a range of reasonable charge states. Matched values near or above the mass threshold for isotopic resolution on our mass spectrometer were written along with their fragment list into *.msalign files to search with TopPIC. Adding additional matching constraints, including 1) a requirement that each match was isolated at least twice within a reasonable retention period, and 2) a requirement that matches must come from scans with no greater than 15 UniDec picked peaks, greatly decreased the number of false matches (**Figure S3**). A set maximum number of picked peaks proved essential as UniDec deconvolution results are output in terms of relative abundance, meaning a noisy spectrum will produce many peaks above the intensity threshold, greatly increasing the likelihood of false matches.

By plotting the precursor mass and charge values across multiple deconvolution methods, we observed significant improvement from the correction strategy over TopFD and FLASHDeconv (**Figure 3A, B**). The presence of vertical clusters of adjacent charge states at the same precursor mass indicates the presence of isotopically unresolved proteins that have been successfully corrected. While TopFD cannot deconvolute isotopically unresolved spectra, FLASHDeconv has been reported as capable of large protein deconvolution.^42^ Interestingly, with our dataset, FLASHDeconv seemed to perform well with high mass, low charge proteins with less congestion in their charge state envelopes, such as the two prominent precursor clusters which fall outside the linear precursor mass to charge relationship. These two clusters, one at approximately 65 kDa (50+) and the other at 80 kDa (40+), correspond to proteins with disulfide bonds that were not fully reduced. However, FLASHDeconv appeared to miss many precursor clusters which were found with the correction algorithm.

**Figure 3.**
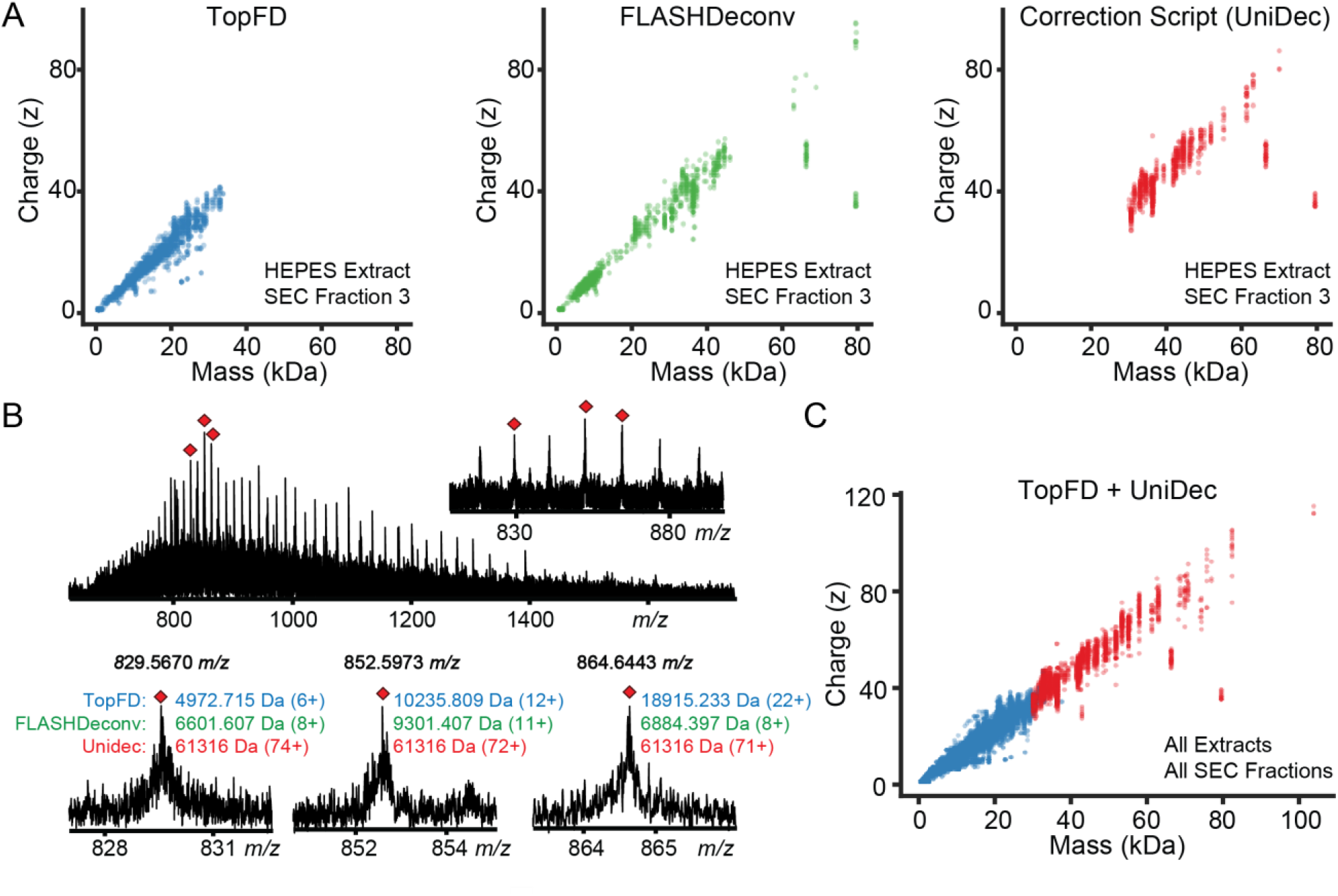
Precursor mass correction and identification of isotopically unresolved proteins. **A)** Precursor mass-charge distributions from a representative data file (HEPES extract, SEC fraction 3 processed using TopFD, FLASHDeconv, or the UniDec-based correction algorithm. **B)** Representative MS1 spectrum and isolated precursors for an isotopically unresolved species illustrating erroneous mass assignments by TopFD and FLASHDevonv and correct assignment by the correction algorithm. **C)** Combined precursor mass-charge distribution across the searched dataset, incorporating original TopFD assignments and UniDec-corrected assignments for unresolved precursors.

The final searched dataset consisted of a set of the original *.msalign files output from TopFD, and a set where only scans with corrected precursor mass values were written. This strategy combines the strengths of TopFD for feature detection and deconvolution of isotopically resolved protein or fragment ions with the strengths of UniDec for unresolved proteins, resulting in a precursor mass distribution from 4 kDa to over 100 kDa (**Figure 3C**). A TopPIC search of the files generated by precursor mass correction resulted in identification of 39 unique proteins over 30 kDa, a greater than tenfold increase over the original TopPIC search. This corresponded to 38 nonredundant unique proteins, totaling to 41 unique proteins >30 kDa when combined with the original TopFD results. The average mass of the unique proteins identified from the precursor-corrected data was 41.6 kDa, while the average mass of unique proteins from the TopFD precursor data was 12.6 kDa. Examples of large proteins identified following precursor mass correction and manually verified by MS/MS analysis are included in **Figure S4**. The largest proteins identified via this strategy were glucose-6-phosphate isomerase (63.1 kDa) and albumin (66.4 kDa).

The corrected precursor masses correspond to the average mass of the proteoform to the nearest Da, whereas TopPIC expects highly accurate monoisotopic mass. Because TopPIC has a generous window for unexpected mass shifts, proteoforms can still be identified using their average precursor mass when sufficient fragmentation evidence is available. In these cases, TopPIC will report an unexpected mass shift in the interior of the sequence uncovered by fragmentation which corresponds to the difference between the expected monoisotopic mass and the average mass. This causes inaccuracy in the calculated E-values and can prevent identification of proteoforms in cases where additional mass shifts are present near a terminus. This also results in an overestimation of unique proteoforms as many proteoforms will be reported by TopPIC with small changes in the location of the unexpected mass shift. Overall, top-down database search tools would benefit from average mass search capabilities and from deconvolution algorithms capable of handling both resolved and unresolved isotopic envelopes in the same data file.

### Identification of Large Proteins by Data-Independent Acquisition and Intact-Mass-Guided Candidate Discovery

Although precursor mass correction increased protein identification coverage above 30 kDa by more than tenfold, many detected large proteins remained unidentified. As protein mass increases, signal is distributed across more charge states and isotopologues, resulting in an exponential decrease in signal-to-noise ratio.^16^ The number of possible terminal and internal fragments also increases with sequence length, distributing fragment-ion signal across more products and increasing spectral complexity. Together, these effects make quality DDA MS/MS spectra difficult to obtain for large proteins on chromatographic timescales. Therefore, we developed a DIA method coupled with intact-mass-guided candidate discovery and MS/MS validation to extend identification to larger proteins. This DIA method consisted of two alternating fragmentation windows of 100 *m/z*, together spanning a mass range from 800-1000 *m/z*, where most charge states belonging to large proteins were detected (**Figure S5**). In contrast to a DDA method which typically selects a single charge state representing only a small fraction of the signal for a large protein, this DIA method provides broader windows covering multiple charge states simultaneously, increasing fragment ion signal and percentage of bond cleavages (**Figure S6**). However, this data could not be analyzed by existing top-down DIA tools such as TopDIA^62^ which requires isotopic resolution to perform precursor-fragment grouping and produce pseudo spectra. We therefore determined chromatographic elution windows for individual large-protein features and summed the corresponding DIA MS/MS spectra for subsequent interpretation. Reduced coelution following online SEC–RPLC facilitated this analysis.

To identify large proteins from the DIA data, we developed an intact-mass-guided strategy to propose reasonable candidates for unidentified spectra and validate their identities (**Figure 4**). First, we generated a protein library based on a bottom-up proteomics (BUP) dataset from human heart tissue published previously by our lab.^43^ Theoretical average masses for each protein in the library were generated based on the protein sequence and a select list of UniProt annotations consisting of signal/transit peptide cleavages, N-terminal Met cleavages, and N-terminal acetylation (**Figure 4B**). This is comparable to the use of UniProt data in ProSight’s “PTM warehouse” or in MetaMorpheus, although less comprehensive to limit the search space to the most likely candidates.^63,64^ Then, a list of detected average matches were matched against the library theoretical masses. We also considered a small list of mass shifts corresponding to various N-terminal states that are commonly missing or differ from UniProt annotations, for example, a 131 Da mass shift corresponding to an unreported Met cleavage (**Table S2**). Candidates were sorted into three groups based on their proximity to the detected mass: 1) “exact matches” within 2 Da, 2) “shift rescues” where an N-terminal variable shift brings the mass within 2 Da, and 3) “vicinity candidates”, the nearest proteins that are not exact mass matches. Candidates in each group were then sorted by their label-free quantification (LFQ) abundance from the bottom-up study, as top-down proteomics is generally limited to high abundance proteins, especially with large proteins. Alternatively, a library consisting of all UniProt human proteins filtered by transcript abundance in heart tissue performed comparably. Plausible candidates were validated against their respective MS/MS spectra in MASH Native (**Figure 4C**), and a putative protein identity was accepted for spectra with sufficient fragmentation evidence.

**Figure 4.**
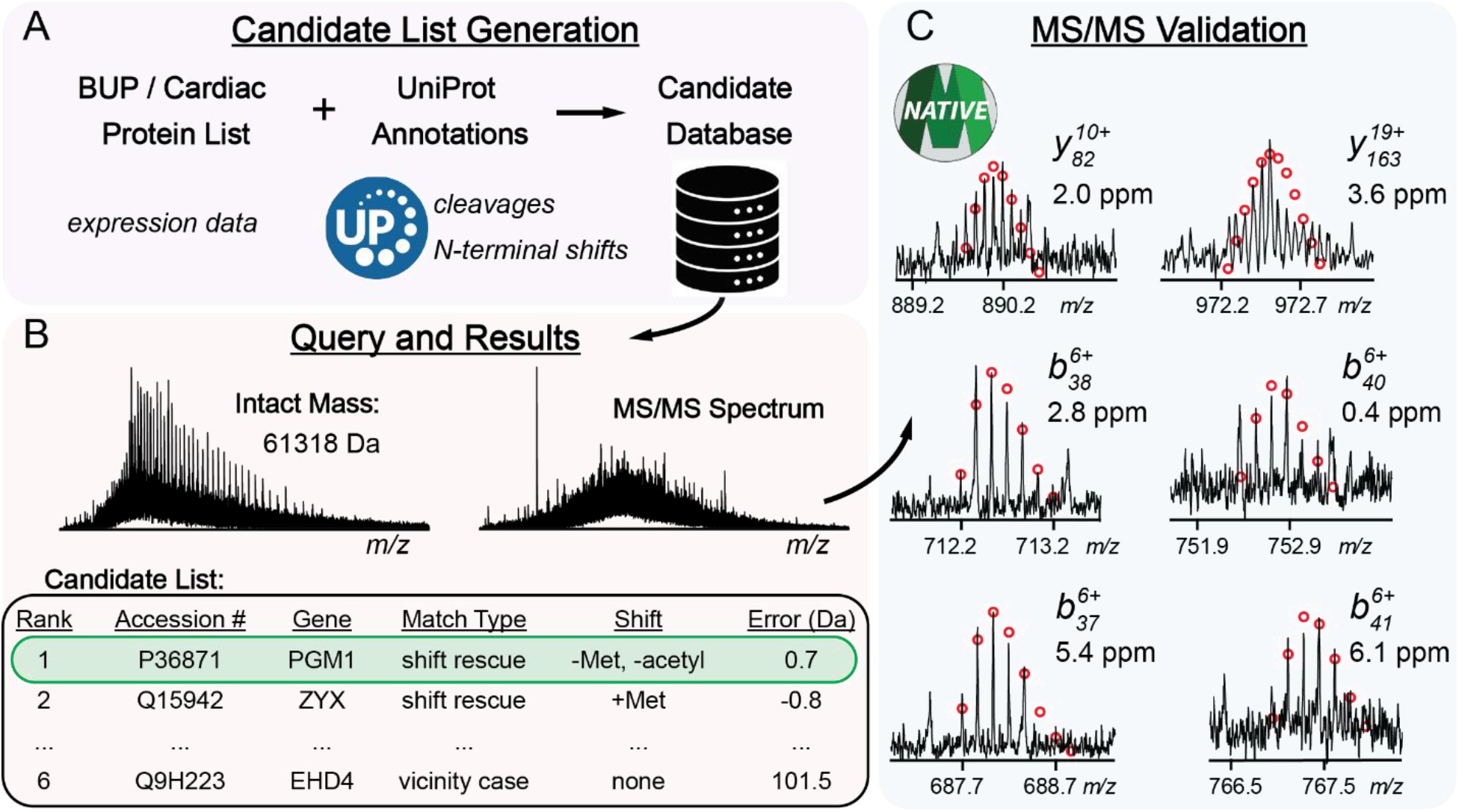
Intact-mass-guided candidate discovery and validation for identification of large proteins. **A)** Construction of a candidate library from proteins previously identified in human heart tissue by bottom-up proteomics (BUP)^43^. Theoretical average masses were calculated using selected UniProt annotations, including signal/transit peptide cleavages, N-terminal methionine removal, and N-terminal acetylation. **B)** Representative intact-mass query and candidate-ranking process. Candidates were grouped as “exact matches” within 2 Da of the measured mass, “shift rescues” brought within 2 Da by an additional N-terminal alteration (**Table S2**), or “vicinity candidates” representing the nearest remaining theoretical masses. Within each group, candidates were ranked by BUP-derived label-free quantification abundance. **C)** Representative MS/MS validation of a candidate identity through spectral deconvolution and terminal fragment-ion matching in MASH Native.

Following this scheme, we identified proteins including alpha-actinin-2 (ACTN2), a 104 kDa Z-disk structural protein,^65^ and the 82.4 kDa mitochondrial aconitate hydratase (ACON) (**Figure 5A**). Based on accurate mass and *b* ion detection, ACTN2 was determined to be N-terminally acetylated, while ACON was determined to have an N-terminal pyro-Glu following transit peptide cleavage. Other identified proteins include vinculin, a 117 kDa actin-binding protein, for which we observed two proteoforms, one corresponding to the Met-cleaved form and another with a +127 Da mass shift. We identified stress-induced-phosphoprotein 1 and desmin, both of which displayed a main peak and secondary +80 Da proteoform which most likely correspond to phosphorylation. These examples and a collection of other identified large proteins are shown in **Figure S7**, along with additional examples of validated MS/MS spectra (**Figures S8-10**). Serotransferrin (TRFE), a 79.6 kDa protein, was identified through a 15-length sequence tag using the *de novo* sequencing pipeline in Omniscape and MS-BLAST search.^45^ Due to an abundance of unreduced disulfide bonds, the charge state distribution of serotransferrin was beyond the range isolated in our DIA method, so we instead used a summed DDA spectra. The intact-mass-guided method was unable to propose serotransferrin as a candidate because it contains a large mass shift of multiple thousand Da from its canonical sequence, likely caused by glycosylation.^66^ After accounting for mass shifts due to a signal peptide cleavage and disulfide bonds near the N-terminus, we were able to validate the series of *b* ions comprising the sequence tag in MASH (**Figure 5B**). Overall, the intact-mass-guided candidate discovery and one sequence-tag analysis extended large-protein coverage to 24 additional protein identities, with an average mass of approximately 73 kDa.

**Figure 5.**
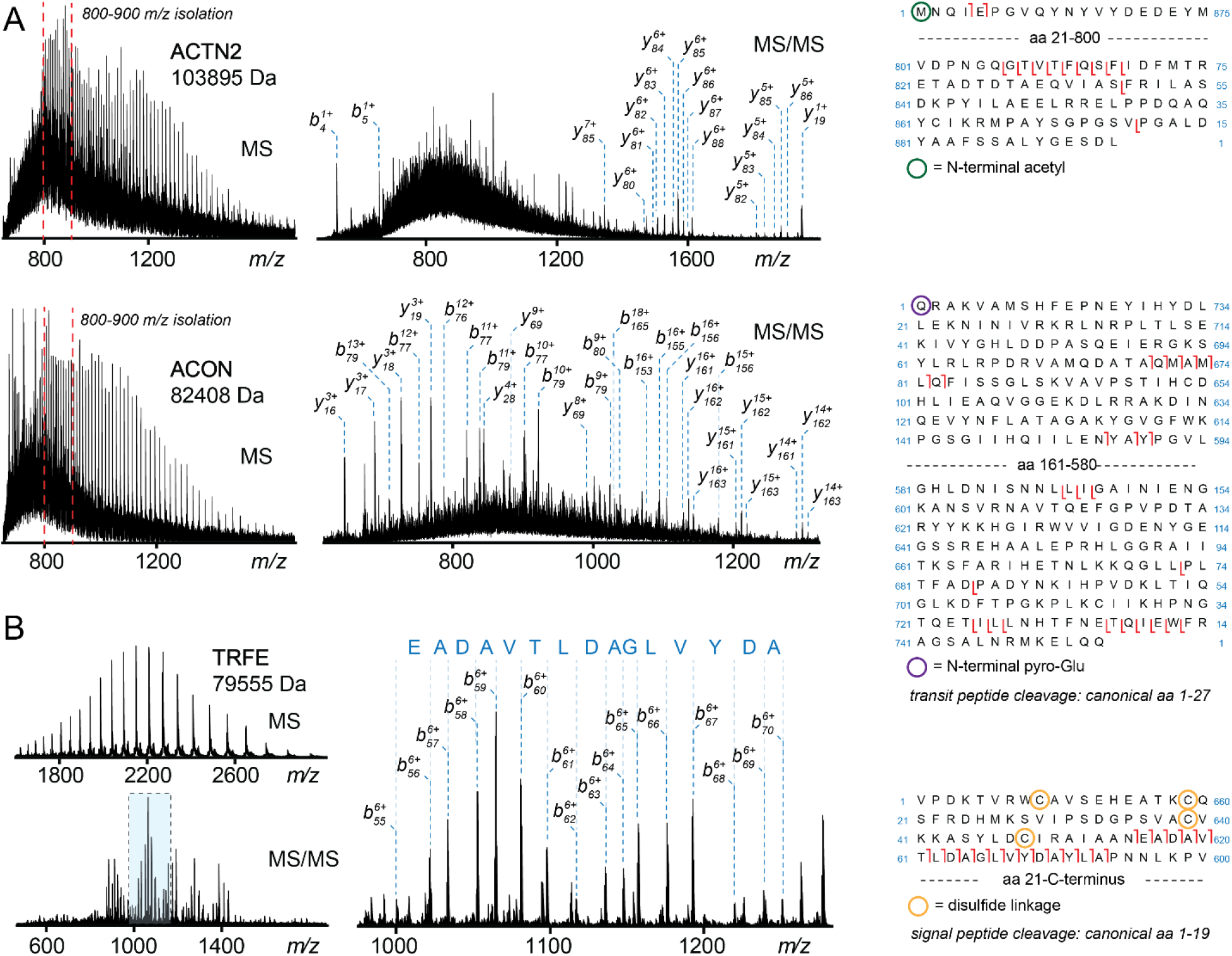
Identification of large proteins using the intact-mass-guided candidate discovery and complementary de novo sequence-tag analysis. **A)** Representative intact mass spectra, annotated MS/MS spectra, and sequence maps for alpha-actinin-2 (ACTN2, 104 kDa) and mitochondrial aconitate hydratase (ACON, 82.4 kDa), identified through intact-mass-guided candidate discovery and MS/MS validation. Intact-mass agreement and N-terminal *b* ions support N-terminal acetylation of ACTN2 and pyroglutamate formation following transit peptide cleavage of ACON. **B)** Identification of serotransferrin (TRFE, 79.6 kDa) using summed DDA spectra. Its observed precursor charge states lay outside the DIA isolation windows, and its intact mass deviation exceeded the shifts accommodated by the TopPIC search and intact-mass-guided candidate discovery settings. A 15-residue sequence tag generated using Bruker OmniScape *de novo* sequencing yielded a candidate identity through an MS-BLAST search. The corresponding *b*-ion series was validated in MASH Native after accounting for signal peptide cleavage and disulfide bonds in the N-terminal region.

### An Integrated Experimental and Informatics Method Extends the Mass Range of Top-Down Proteomics

By integrating online SEC–RPLC separation, precursor mass correction for conventional DDA database searching, and DIA coupled with intact-mass-guided candidate discovery and MS/MS validation, we significantly improved the detection and identification of large proteins. Overall, this integrated identified 703 nonredundant proteoforms corresponding to 236 unique proteins from human heart tissue. A STRING analysis of the set of genes corresponding to all unique proteins using a Leiden algorithm (MCL) resolution parameter of 1.0 produced 4 clusters: cytoskeleton in muscle cells/contractile muscle fiber, carbon metabolism/aerobic respiration, oxidative phosphorylation, and peptide chain elongation (**Figure 6A**). Enrichment of these subproteomes aligns with previous experiments utilizing the same three-step extraction method on human heart tissue.^37^ These integrated identification strategies progressively expanded large-protein coverage. The original TopFD/TopPIC workflow identified only 3 proteins >30 kDa. Precursor mass correction step resulted in an additional 38 nonredundant unique proteins >30 kDa. The intact-mass-guided candidate discovery workflow resulted in an additional 24 nonredundant unique proteins >30 kDa. In total, this corresponds to 65 proteins >30 kDa, 48 proteins >40 kDa, and 22 proteins >50 kDa (**Figure 6B-E)**. The largest proteins identified were myosin-7 (223 kDa), vinculin (117 kDa), and alpha-actinin-2 (104 kDa).

**Figure 6.**
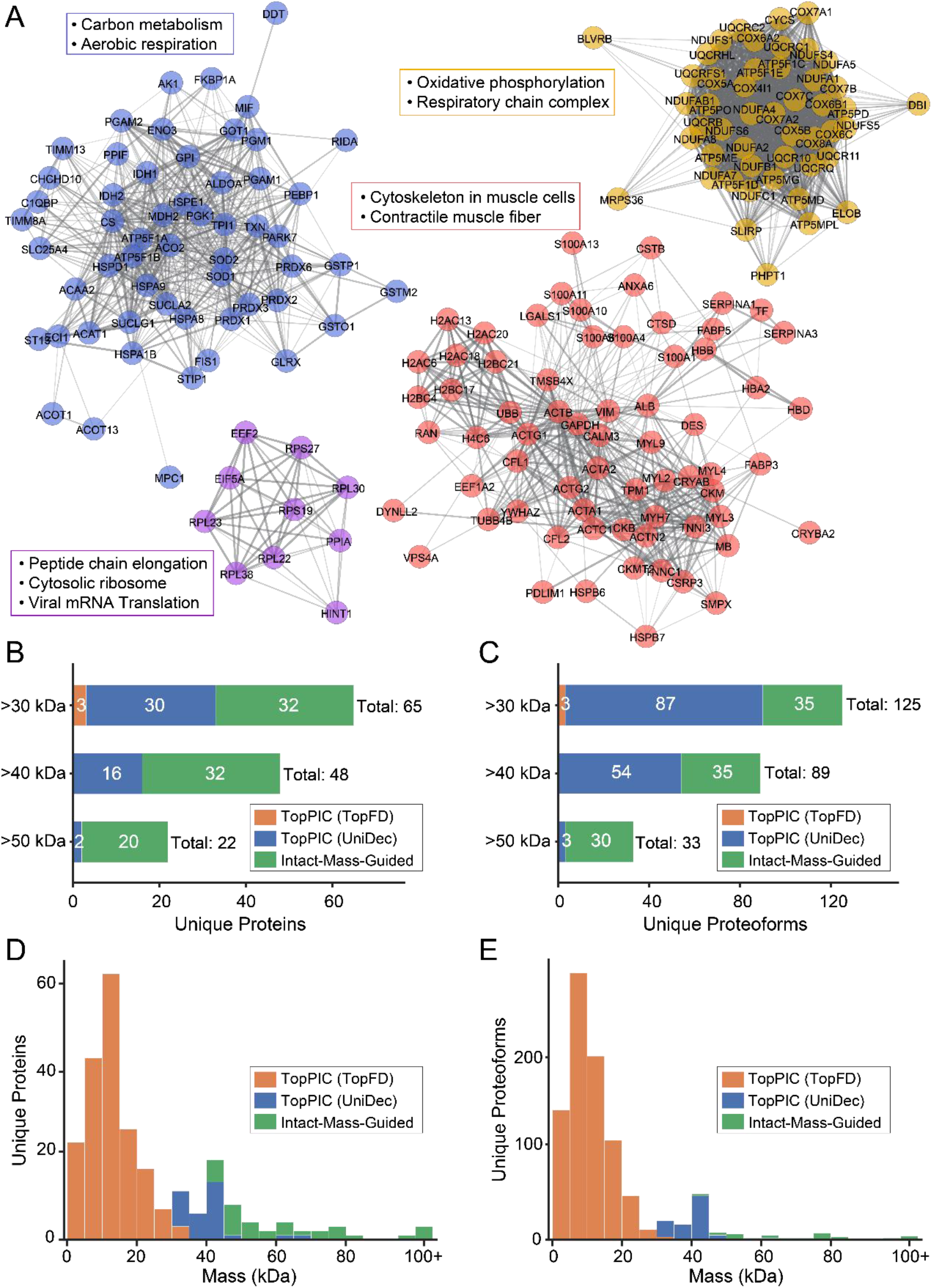
Summary of all proteins identified from human heart tissue using the integrated experimental and informatics workflow. **A)** STRING diagram constructed from all identified proteins. Clustering was performed using a Leiden inflation resolution parameter of 1.0, resulting in 4 clusters. **B-C)** Bar plots displaying the number of unique identified proteins and proteoforms above select mass thresholds (30, 40, and 50 kDa) according to identification method. **D-E)** Histogram plotting the molecular weight of unique proteins and proteoforms according to identification method.

Despite advances in separations, discovery top-down proteomics datasets remain dominated by small proteoforms due to the exponential decrease of signal-to-noise ratio with increasing molecular weight.^16^ For example, a previous online 2D-LC study conducted by the Wu lab coupling high pH and low pH RPLC reported an average proteoform mass of 7.3 kDa.^18^ Compared to other chromatographic modes, SEC greatly extends the detectable molecular weight range of proteoforms by reducing coelution with smaller proteins that contribute to spectral interference and ion suppression of larger proteins. Here, our online 2D-LC (SEC-RPC) method resulted in an average proteoform mass of 15.8 kDa, and an average unique protein mass of 23.4 kDa.

However, extending the detectable mass range through size-based separation alone does not necessarily translate into identification of large proteins. For example, a previous top-down study conducted by the Sun lab combining SEC, RPLC, and capillary zone electrophoresis (CZE) identified roughly 23,000 proteoforms from over 400 raw files, yet the identified average proteoform mass was approximately 6.2 kDa, and no proteoform identifications were over ∼26 kDa.^19^ Similarly, 93% of the nearly 30,000 proteoforms in the Blood Proteoform Atlas were below 20 kDa despite GELFrEE fractionation.^17^ These studies demonstrate extensive proteoform coverage, highlighting the persistent challenge of large-protein identification.

Our previous study using offline sSEC-RPLC for top-down proteomics of human heart tissue reported detection of large proteins by MS1, but targeted MS/MS identification was demonstrated for just two isotopically resolved proteins between 40-50 kDa. Subsequent analysis using Proteoform Suite generated candidates from intact average masses, but with an extremely high FDR for large proteins (82.7% FDR above 50 kDa), and only 3 unique proteins >50 kDa were identified by targeted MS/MS.^36^ Thus, SEC enables detection, but not necessarily identification of large proteins. Here, our strategy integrating online SEC–RPLC with precursor mass correction and intact-mass-guided candidate discovery allowed us to identify 48 unique proteins >40 kDa and 22 unique proteins >50 kDa with a maximum identified mas of 223 kDa.

Additionally, the online 2D-LC (SEC–RPLC) approach streamlines sample transfer by eliminating offline fraction collection and reinjection, reducing manual handling and minimizing sample loss. This study used a maximum of 25 μg protein per injection, compared with approximately 300 μg in our previous offline sSEC–RPLC study^27^, representing a significant improvement in sensitivity. Sensitivity could be improved further by addition of sample storage loops or other interface to allow a ^2^D analysis of every ^1^D fraction from one sample injection, enhancing sample utilization and automation.

Although the integrated strategy expands identification of large proteoforms, further development is needed to automate data processing and incorporate these approaches into routine discovery top-down proteomics workflows. An important next step is to integrate deconvolution methods capable of processing both isotopically resolved and unresolved precursor ions within the same data file. Recently developed deconvolution algorithms such as kDecon from the Prosight team are moving toward this capability.^67^ Denatured top-down proteomics searching tools such as TopPIC and others would benefit from the capability to identify proteoforms from both monoisotopic and average precursor masses with proper E-value calculation and FDR control. Another possibility would be to extend the capabilities of recent DIA strategies such as TopDIA^62^ to associate fragment ions with isotopically unresolved precursor masses. A recent full-window DIA processing method has shown some potential of integrating multiple deconvolution methods, although it did not specifically address isotopically unresolved proteins.^68^

## Conclusions

This study represents a multifaceted approach to extending the mass range of top-down proteomics. We have developed an integrated experimental and informatics strategy to enable identification of large proteins, including online SEC-RPLC-MS/MS, automated precursor mass correction, DIA methods for large protein fragmentation, and an intact-mass-guided candidate discovery pipeline. Online SEC–RPLC-MS/MS facilitates separation and detection of large proteoforms. Automated precursor mass correction enables conventional database searching of isotopically unresolved precursors. For species that remain unidentified, DIA tailored to large proteins enhances fragment ion signal for intact-mass-guided candidate discovery and MS/MS validation, complemented by *de novo* sequence-tag analysis. Applied to human heart tissue, precursor mass correction increases the number of identified proteins >30 kDa from 3 to 41, while DIA combined with the intact-mass-guided pipeline adds 24 unique proteins averaging approximately 73 kDa. Utilizing this integrated strategy, we identified 703 proteoforms from 236 unique proteins including 65 unique proteins >30 kDa from an online 2D-LC-MS/MS dataset from human heart tissue. Of these unique proteins, 22 were greater than 50 kDa with identified masses extending to 223 kDa. We envision that continued development and implementation of similar strategies will substantially expand the coverage of large proteoforms in discovery top-down proteomics.

## Acknowledgements

The authors thank Christopher J. Wike of PolyLC for providing chromatographic columns. We thank Dr. Detlev Suckau and Dr. Guillaume Tremintin of Bruker Daltonics for providing use of the Omniscape software. We also express gratitude to the donor and their loved ones for their generous tissue donation. We thank James Anderson and Erin Halpin at the University of Wisconsin Organ and Tissue Donation for coordination of donor heart collection. Support for this research was provided by the University of Wisconsin– Madison Office of the Vice Chancellor for Research with funding from the Wisconsin Alumni Research Foundation (to Y.G.). Additional support for this research was provided by NIH 5R01HL156670-05 (to S.D.M and C.C.S.). H.-J. C. acknowledges support from the American Heart Association (AHA) under Award Number 26PRE1561668. Some figure components are assets from BioRender.com. The authors declare the following competing financial interest: Ying Ge is a co-inventor on a patent that covers the detergent Azo (US Patent No: US11,567,085 B2).

## Supporting Information

Additional Tables and Figures as noted in the text. Collisional energy table used for autoMS/MS (Table S1). Mass shifts considered for candidate selection (Table S2). Values used to calculate 2D-LC peak capacity (Table S3). Valve configuration for online SEC-RPLC-MS/MS (Figure S1). Representative 2D-LC-MS data from the HEPES extraction (Figure S2). Improving the fidelity of the precursor correction algorithm (Figure S3). Representative large proteins identified by TopPIC following precursor mass correction (Figure S4). Representative DIA MS/MS spectra for a 63 kDa protein (Figure S5). Comparison of DDA and DIA MS/MS spectra (Figure S6). Representative large proteins identified using the candidate discovery algorithm and MS/MS validation (Figure S7). Identification of NADH-ubiquinone oxidoreductase 75 kDa subunit, mitochondrial (NDUS1, 77 kDa) (Figure S8). Identification of ATP synthase F(1) complex subunit alpha, mitochondrial (ATPA, 55 kDa) (Figure S9). Identification of pyruvate kinase M1 (PKM1, 58 kDa) (Figure S10). (PDF)

## References

(1) The Consortium for Top Down Proteomics; Smith, L. M.; Kelleher, N. L. Proteoform: A Single Term Describing Protein Complexity. Nat Methods 2013, 10 (3), 186–187. 10.1038/nmeth.2369

(2) Smith, L. M.; Agar, J. N.; Chamot-Rooke, J.; Danis, P. O.; Ge, Y.; Loo, J. A.; Paša-Tolić, L.; Tsybin, Y. O.; Kelleher, N. L.; The Consortium for Top-Down Proteomics. The Human Proteoform Project: Defining the Human Proteome. Sci. Adv. 2021, 7 (46), eabk0734. 10.1126/sciadv.abk0734

(3) Smith, L. M.; Kelleher, N. L. Proteoforms as the next Proteomics Currency. Science 2018, 359 (6380), 1106–1107. 10.1126/science.aat1884

(4) Roberts, D. S.; Loo, J. A.; Tsybin, Y. O.; Liu, X.; Wu, S.; Chamot-Rooke, J.; Agar, J. N.; Paša-Tolić, L.; Smith, L. M.; Ge, Y. Top-down Proteomics. Nat Rev Methods Primers 2024, 4 (1), 38. 10.1038/s43586-024-00318-2

(5) Po, A.; Eyers, C. E. Top-Down Proteomics and the Challenges of True Proteoform Characterization. J. Proteome Res. 2023, 22 (12), 3663–3675. 10.1021/acs.jproteome.3c00416

(6) Kaulich, P. T.; Tholey, A. Top-Down Proteomics: Why and When? Proteomics 2025, 25 (24), 6–12. 10.1002/pmic.202400338

(7) Brodbelt, J. S. Deciphering Combinatorial Post-Translational Modifications by Top-down Mass Spectrometry. Current Opinion in Chemical Biology 2022, 70, 102180. 10.1016/j.cbpa.2022.102180

(8) Rogers, H. T.; Gregorich, Z. R.; Gant, M. S.; Gao, Z.; Chamot-Rooke, J.; Ge, Y. Proteoforms in Disease: Biomedical Applications of Top-Down Proteomics. Mass Spectrometry Reviews 2026, mas.70042. 10.1002/mas.70042

(9) Brown, K. A.; Melby, J. A.; Roberts, D. S.; Ge, Y. Top-down Proteomics: Challenges, Innovations, and Applications in Basic and Clinical Research. Expert Review of Proteomics 2020, 17 (10), 719–733. 10.1080/14789450.2020.1855982

(10) Nickerson, J. L.; Baghalabadi, V.; Rajendran, S. R. C. K.; Jakubec, P. J.; Said, H.; McMillen, T. S.; Dang, Z.; Doucette, A. A. Recent Advances in Top-down Proteome Sample Processing Ahead of MS Analysis. Mass Spectrometry Reviews 2023, 42 (2), 457–495. 10.1002/mas.21706

(11) Melby, J. A.; Roberts, D. S.; Larson, E. J.; Brown, K. A.; Bayne, E. F.; Jin, S.; Ge, Y. Novel Strategies to Address the Challenges in Top-Down Proteomics. J. Am. Soc. Mass Spectrom. 2021, 32 (6), 1278–1294. 10.1021/jasms.1c00099

(12) Walker, J. N.; Brodbelt, J. S. Ion Activation Methods for Top-Down Proteomics. Mass Spectrometry Reviews 2026, mas.70023. 10.1002/mas.70023

(13) Cupp-Sutton, K. A.; Wu, S. High-Throughput Quantitative Top-down Proteomics. Molecular Omics 2020, 16 (2), 91–99. 10.1039/C9MO00154A

(14) Zhu, G.; Alvarez Rodriguez, D. A.; Gould, N.; Ivanov, A. R.; Jooß, K.; Sun, L. Recent Advances (2023–2025) of Capillary Electrophoresis-Mass Spectrometry (CE-MS) for Top-Down Proteomics. Mass Spectrometry Reviews 2026, mas.70028. 10.1002/mas.70028

(15) Fornelli, L.; Toby, T. K. Characterization of Large Intact Protein Ions by Mass Spectrometry: What Directions Should We Follow? Biochimica et Biophysica Acta (BBA) – Proteins and Proteomics 2022, 1870 (4), 140758. 10.1016/j.bbapap.2022.140758

(16) Compton, P. D.; Zamdborg, L.; Thomas, P. M.; Kelleher, N. L. On the Scalability and Requirements of Whole Protein Mass Spectrometry. Anal. Chem. 2011, 83 (17), 6868–6874. 10.1021/ac2010795

(17) Melani, R. D.; Gerbasi, V. R.; Anderson, L. C.; Sikora, J. W.; Toby, T. K.; Hutton, J. E.; Butcher, D. S.; Negrão, F.; Seckler, H. S.; Srzentić, K.; Fornelli, L.; Camarillo, J. M.; LeDuc, R. D.; Cesnik, A. J.; Lundberg, E.; Greer, J. B.; Fellers, R. T.; Robey, M. T.; DeHart, C. J.; Forte, E.; Hendrickson, C. L.; Abbatiello, S. E.; Thomas, P. M.; Kokaji, A. I.; Levitsky, J.; Kelleher, N. L. The Blood Proteoform Atlas: A Reference Map of Proteoforms in Human Hematopoietic Cells. Science 2022, 375 (6579), 411–418. 10.1126/science.aaz5284

(18) Wang, Z.; Yu, D.; Cupp-Sutton, K. A.; Liu, X.; Smith, K.; Wu, S. Development of an Online 2D Ultrahigh-Pressure Nano-LCSystem for High-pH and Low-pH Reversed Phase Separation in Top-DownProteomics. Analytical Chemistry 2020, 92 (19), 12774–12777. 10.1021/acs.analchem.0c03395

(19) McCool, E. N.; Xu, T.; Chen, W.; Beller, N. C.; Nolan, S. M.; Hummon, A. B.; Liu, X.; Sun, L. Deep Top-down Proteomics Revealed Significant Proteoform-Level Differences between Metastatic and Nonmetastatic Colorectal Cancer Cells. Sci. Adv. 2022, 8 (51), eabq6348. 10.1126/sciadv.abq6348

(20) Catherman, A. D.; Durbin, K. R.; Ahlf, D. R.; Early, B. P.; Fellers, R. T.; Tran, J. C.; Thomas, P. M.; Kelleher, N. L. Large-Scale Top-down Proteomics of the Human Proteome: Membrane Proteins, Mitochondria, and Senescence. Molecular & Cellular Proteomics 2013, 12 (12), 3465–3473. 10.1074/mcp.M113.030114

(21) Xu, T.; Wang, Q.; Wang, Q.; Sun, L. Mass Spectrometry-Intensive Top-down Proteomics: An Update on Technology Advancements and Biomedical Applications. Anal. Methods 2024, 16 (28), 4664–4682. 10.1039/D4AY00651H

(22) Forte, E.; Sanders, J. M.; Pla, I.; Kanchustambham, V. L.; Hollas, M. A. R.; Huang, C.-F.; Sanchez, A.; Peterson, K. N.; Melani, R. D.; Huang, A.; Polineni, P.; Doll, J. M.; Dietch, Z.; Kelleher, N. L.; Ladner, D. P. Top-Down Proteomics Identifies Plasma Proteoform Signatures of Liver Cirrhosis Progression. Molecular & Cellular Proteomics 2024, 23 (12), 100876. 10.1016/j.mcpro.2024.100876

(23) Kaulich, P. T.; Jeong, K.; Kohlbacher, O.; Tholey, A. Influence of Different Sample Preparation Approaches on Proteoform Identification by Top-down Proteomics. Nat Methods 2024, 21 (12), 2397–2407. 10.1038/s41592-024-02481-6

(24) Durbin, K. R.; Fornelli, L.; Fellers, R. T.; Doubleday, P. F.; Narita, M.; Kelleher, N. L. Quantitation and Identification of Thousands of Human Proteoforms below 30 kDa. J. Proteome Res. 2016, 15 (3), 976–982. 10.1021/acs.jproteome.5b00997

(25) Cheon, D. H.; Nam, E. J.; Park, K. H.; Woo, S. J.; Lee, H. J.; Kim, H. C.; Yang, E. G.; Lee, C.; Lee, J. E. Comprehensive Analysis of Low-Molecular-Weight Human Plasma Proteome Using Top-Down Mass Spectrometry. J. Proteome Res. 2016, 15 (1), 229–244. 10.1021/acs.jproteome.5b00773

(26) The UniProt Consortium; Bateman, A.; Martin, M.-J.; Orchard, S.; Magrane, M.; Adesina, A.; Ahmad, S.; Bowler-Barnett, E. H.; Bye-A-Jee, H.; Carpentier, D.; Denny, P.; Fan, J.; Garmiri, P.; Gonzales, L. J. D. C.; Hussein, A.; Ignatchenko, A.; Insana, G.; Ishtiaq, R.; Joshi, V.; Jyothi, D.; Kandasaamy, S.; Lock, A.; Luciani, A.; Luo, J.; Lussi, Y.; Marin, J. S. M.; Raposo, P.; Rice, D. L.; Santos, R.; Speretta, E.; Stephenson, J.; Totoo, P.; Tyagi, N.; Urakova, N.; Vasudev, P.; Warner, K.; Wijerathne, S.; Yu, C. W.-H.; Zaru, R.; Bridge, A. J.; Aimo, L.; Argoud-Puy, G.; Auchincloss, A. H.; Axelsen, K. B.; Bansal, P.; Baratin, D.; Batista Neto, T. M.; Blatter, M.-C.; Bolleman, J. T.; Boutet, E.; Breuza, L.; Gil, B. C.; Casals-Casas, C.; Echioukh, K. C.; Coudert, E.; Cuche, B.; De Castro, E.; Estreicher, A.; Famiglietti, M. L.; Feuermann, M.; Gasteiger, E.; Gaudet, P.; Gehant, S.; Gerritsen, V.; Gos, A.; Gruaz, N.; Hulo, C.; Hyka-Nouspikel, N.; Jungo, F.; Kerhornou, A.; Mercier, P. L.; Lieberherr, D.; Masson, P.; Morgat, A.; Paesano, S.; Pedruzzi, I.; Pilbout, S.; Pourcel, L.; Poux, S.; Pozzato, M.; Pruess, M.; Redaschi, N.; Rivoire, C.; Sigrist, C. J. A.; Sonesson, K.; Sundaram, S.; Sveshnikova, A.; Wu, C. H.; Arighi, C. N.; Chen, C.; Chen, Y.; Huang, H.; Laiho, K.; Lehvaslaiho, M.; McGarvey, P.; Natale, D. A.; Ross, K.; Vinayaka, C. R.; Wang, Y.; Zhang, J. UniProt: The Universal Protein Knowledgebase in 2025. Nucleic Acids Research 2025, 53 (D1), D609–D617. 10.1093/nar/gkae1010

(27) Cai, W.; Tucholski, T.; Chen, B.; Alpert, A. J.; McIlwain, S.; Kohmoto, T.; Jin, S.; Ge, Y. Top-Down Proteomics of Large Proteins up to 223 kDa Enabled by Serial Size Exclusion Chromatography Strategy. Anal. Chem. 2017, 89 (10), 5467–5475. 10.1021/acs.analchem.7b00380

(28) Han, X.; Jin, M.; Breuker, K.; McLafferty, F. W. Extending Top-Down Mass Spectrometry to Proteins with Masses Greater Than 200 Kilodaltons. Science 2006, 314 (5796), 109–112. 10.1126/science.1128868

(29) Chapman, E. A.; Roberts, D. S.; Tiambeng, T. N.; Andrews, J.; Wang, M.-D.; Reasoner, E. A.; Melby, J. A.; Li, B. H.; Kim, D.; Alpert, A. J.; Jin, S.; Ge, Y. Structure and Dynamics of Endogenous Cardiac Troponin Complex in Human Heart Tissue Captured by Native Nanoproteomics. Nat Commun 2023, 14 (1), 8400. 10.1038/s41467-023-43321-z

(30) Li, H.; Nguyen, H. H.; Ogorzalek Loo, R. R.; Campuzano, I. D. G.; Loo, J. A. An Integrated Native Mass Spectrometry and Top-down Proteomics Method That Connects Sequence to Structure and Function of Macromolecular Complexes. Nature Chem 2018, 10 (2), 139–148. 10.1038/nchem.2908

(31) Liu, X.; Segar, M. W.; Li, S. C.; Kim, S. Spectral Probabilities of Top-down Tandem Mass Spectra. BMC Genomics 2014, 15 (S1), S9. 10.1186/1471-2164-15-S1-S9

(32) Tabb, D. L.; Jeong, K.; Druart, K.; Gant, M. S.; Brown, K. A.; Nicora, C.; Zhou, M.; Couvillion, S.; Nakayasu, E.; Williams, J. E.; Peterson, H. K.; McGuire, M. K.; McGuire, M. A.; Metz, T. O.; Chamot-Rooke, J. Comparing Top-Down Proteoform Identification: Deconvolution, PrSM Overlap, and PTM Detection. J. Proteome Res. 2023, 22 (7), 2199–2217. 10.1021/acs.jproteome.2c00673

(33) Kou, Q.; Xun, L.; Liu, X. TopPIC: A Software Tool for Top-down Mass Spectrometry-Based Proteoform Identification and Characterization. Bioinformatics 2016, 32 (22), 3495–3497. 10.1093/bioinformatics/btw398

(34) Sun, R.-X.; Luo, L.; Wu, L.; Wang, R.-M.; Zeng, W.-F.; Chi, H.; Liu, C.; He, S.-M. pTop 1.0: A High-Accuracy and High-Efficiency Search Engine for Intact Protein Identification. Anal. Chem. 2016, 88 (6), 3082–3090. 10.1021/acs.analchem.5b03963

(35) Park, J.; Piehowski, P. D.; Wilkins, C.; Zhou, M.; Mendoza, J.; Fujimoto, G. M.; Gibbons, B. C.; Shaw, J. B.; Shen, Y.; Shukla, A. K.; Moore, R. J.; Liu, T.; Petyuk, V. A.; Tolić, N.; Paša-Tolić, L.; Smith, R. D.; Payne, S. H.; Kim, S. Informed-Proteomics: Open-Source Software Package for Top-down Proteomics. Nat Methods 2017, 14 (9), 909–914. 10.1038/nmeth.4388

(36) Schaffer, L. V.; Tucholski, T.; Shortreed, M. R.; Ge, Y.; Smith, L. M. Intact-Mass Analysis Facilitating the Identification of Large Human Heart Proteoforms. Anal. Chem. 2019, 91 (17), 10937–10942. 10.1021/acs.analchem.9b02343

(37) Gao, Z.; Rossler, K. J.; Rogers, H. T.; Gregorich, Z. R.; Aballo, T. J.; Chapman, E. A.; Price, S. J.; Hermsen, J.; De Lange, W. J.; Zhu, Y.; Zhang, B.; Chen, J.; Ralphe, J. C.; Kamp, T. J.; Ge, Y. Global Proteoform Alterations Across Multiple Cellular Compartments Underlie Obstructive Hypertrophic Cardiomyopathy. Circ: Heart Failure 2025, 18 (12). 10.1161/CIRCHEARTFAILURE.125.012899

(38) Rogers, H. T.; Roberts, D. S.; Larson, E. J.; Melby, J. A.; Rossler, K. J.; Carr, A. V.; Brown, K. A.; Ge, Y. Comprehensive Characterization of Endogenous Phospholamban Proteoforms Enabled by Photocleavable Surfactant and Top-down Proteomics. Anal. Chem. 2023, 95 (35), 13091–13100. 10.1021/acs.analchem.3c01618

(39) Chambers, M. C.; Maclean, B.; Burke, R.; Amodei, D.; Ruderman, D. L.; Neumann, S.; Gatto, L.; Fischer, B.; Pratt, B.; Egertson, J.; Hoff, K.; Kessner, D.; Tasman, N.; Shulman, N.; Frewen, B.; Baker, T. A.; Brusniak, M.-Y.; Paulse, C.; Creasy, D.; Flashner, L.; Kani, K.; Moulding, C.; Seymour, S. L.; Nuwaysir, L. M.; Lefebvre, B.; Kuhlmann, F.; Roark, J.; Rainer, P.; Detlev, S.; Hemenway, T.; Huhmer, A.; Langridge, J.; Connolly, B.; Chadick, T.; Holly, K.; Eckels, J.; Deutsch, E. W.; Moritz, R. L.; Katz, J. E.; Agus, D. B.; MacCoss, M.; Tabb, D. L.; Mallick, P. A Cross-Platform Toolkit for Mass Spectrometry and Proteomics. Nat Biotechnol 2012, 30 (10), 918–920. 10.1038/nbt.2377

(40) Basharat, A. R.; Zang, Y.; Sun, L.; Liu, X. TopFD: A Proteoform Feature Detection Tool for Top–Down Proteomics. Anal. Chem. 2023, 95 (21), 8189–8196. 10.1021/acs.analchem.2c05244

(41) Marty, M. T.; Baldwin, A. J.; Marklund, E. G.; Hochberg, G. K. A.; Benesch, J. L. P.; Robinson, C. V. Bayesian Deconvolution of Mass and Ion Mobility Spectra: From Binary Interactions to Polydisperse Ensembles. Anal. Chem. 2015, 87 (8), 4370–4376. 10.1021/acs.analchem.5b00140

(42) Jeong, K.; Kim, J.; Gaikwad, M.; Hidayah, S. N.; Heikaus, L.; Schlüter, H.; Kohlbacher, O. FLASHDeconv: Ultrafast, High-Quality Feature Deconvolution for Top-Down Proteomics. Cell Systems 2020, 10 (2), 213–218.e6. 10.1016/j.cels.2020.01.003

(43) Aballo, T. J.; Roberts, D. S.; Melby, J. A.; Buck, K. M.; Brown, K. A.; Ge, Y. Ultrafast and Reproducible Proteomics from Small Amounts of Heart Tissue Enabled by Azo and timsTOF Pro. J. Proteome Res. 2021, 20 (8), 4203–4211. 10.1021/acs.jproteome.1c00446

(44) Larson, E. J.; Pergande, M. R.; Moss, M. E.; Rossler, K. J.; Wenger, R. K.; Krichel, B.; Josyer, H.; Melby, J. A.; Roberts, D. S.; Pike, K.; Shi, Z.; Chan, H.-J.; Knight, B.; Rogers, H. T.; Brown, K. A.; Ong, I. M.; Jeong, K.; Marty, M. T.; McIlwain, S. J.; Ge, Y. MASH Native: A Unified Solution for Native Top-down Proteomics Data Processing. Bioinformatics 2023, 39 (6), btad359. 10.1093/bioinformatics/btad359

(45) Shevchenko, A.; Sunyaev, S.; Loboda, A.; Shevchenko, A.; Bork, P.; Ens, W.; Standing, K. G. Charting the Proteomes of Organisms with Unsequenced Genomes by MALDI-Quadrupole Time-of-Flight Mass Spectrometry and BLAST Homology Searching. Anal. Chem. 2001, 73 (9), 1917–1926. 10.1021/ac0013709

(46) Xiu, L.; Valeja, S. G.; Alpert, A. J.; Jin, S.; Ge, Y. Effective Protein Separation by Coupling Hydrophobic Interaction and Reverse Phase Chromatography for Top-down Proteomics. Anal. Chem. 2014, 86 (15), 7899–7906. 10.1021/ac501836k

(47) Valeja, S. G.; Xiu, L.; Gregorich, Z. R.; Guner, H.; Jin, S.; Ge, Y. Three Dimensional Liquid Chromatography Coupling Ion Exchange Chromatography/Hydrophobic Interaction Chromatography/Reverse Phase Chromatography for Effective Protein Separation in Top-Down Proteomics. Anal. Chem. 2015, 87 (10), 5363–5371. 10.1021/acs.analchem.5b00657

(48) Guo, Y.; Cupp-Sutton, K. A.; Zhao, Z.; Anjum, S.; Wu, S. Multidimensional Separations in Top–down Proteomics. Analytical Science Advances 2023, 4 (5–6), 181–203. 10.1002/ansa.202300016

(49) Brown, K. A.; Tucholski, T.; Alpert, A. J.; Eken, C.; Wesemann, L.; Kyrvasilis, A.; Jin, S.; Ge, Y. Top-Down Proteomics of Endogenous Membrane Proteins Enabled by Cloud Point Enrichment and Multidimensional Liquid Chromatography–Mass Spectrometry. Anal. Chem. 2020, 92 (24), 15726–15735. 10.1021/acs.analchem.0c02533

(50) Wang, Z.; Ma, H.; Smith, K.; Wu, S. Two-Dimensional Separation Using High-pH and Low-pH Reversed Phase Liquid Chromatography for Top-down Proteomics. International Journal of Mass Spectrometry 2018, 427, 43–51. 10.1016/j.ijms.2017.09.001

(51) Vellaichamy, A.; Tran, J. C.; Catherman, A. D.; Lee, J. E.; Kellie, J. F.; Sweet, S. M. M.; Zamdborg, L.; Thomas, P. M.; Ahlf, D. R.; Durbin, K. R.; Valaskovic, G. A.; Kelleher, N. L. Size-Sorting Combined with Improved Nanocapillary Liquid Chromatography−Mass Spectrometry for Identification of Intact Proteins up to 80 kDa. Anal. Chem. 2010, 82 (4), 1234–1244. 10.1021/ac9021083

(52) Tian, Z.; Zhao, R.; Tolić, N.; Moore, R. J.; Stenoien, D. L.; Robinson, E. W.; Smith, R. D.; Paša-Tolić, L. Two-dimensional Liquid Chromatography System for Online Top-down Mass Spectrometry. Proteomics 2010, 10 (20), 3610–3620. 10.1002/pmic.201000367

(53) Baghdady, Y. Z.; Schug, K. A. Online Comprehensive High pH Reversed Phase × Low pH Reversed Phase Approach for Two-Dimensional Separations of Intact Proteins in Top-Down Proteomics. Anal. Chem. 2019, 91 (17), 11085–11091. 10.1021/acs.analchem.9b01665

(54) Stoll, D. R.; Carr, P. W. Two-Dimensional Liquid Chromatography: A State of the Art Tutorial. Anal. Chem. 2017, 89 (1), 519–531. 10.1021/acs.analchem.6b03506

(55) Fischer, M. S.; Rogers, H. T.; Chapman, E. A.; Jin, S.; Ge, Y. Native Top-Down Proteomics of Endogenous Protein Complexes Enabled by Online Two-Dimensional Liquid Chromatography. Anal. Chem. 2025, 97 (25), 13663–13671. 10.1021/acs.analchem.5c02341

(56) Yau, W. W.; Ginnard, C. R.; Kirkland, J. J. Broad-Range Linear Calibration in High-Performance Size-Exclusion Chromatography Using Column Packings with Bimodal Pores. Journal of Chromatography A 1978, 149, 465–487. 10.1016/S0021-9673(00)81005-5

(57) Gritti, F.; Fekete, S. Band Broadening in Pore Size Gradient Size Exclusion Chromatography Columns. Journal of Chromatography A 2026, 1765, 466523. 10.1016/j.chroma.2025.466523

(58) Brown, K. A.; Chen, B.; Guardado-Alvarez, T. M.; Lin, Z.; Hwang, L.; Ayaz-Guner, S.; Jin, S.; Ge, Y. A Photocleavable Surfactant for Top-down Proteomics. Nat Methods 2019, 16 (5), 417–420. 10.1038/s41592-019-0391-1

(59) Davis, J. M.; Stoll, D. R.; Carr, P. W. Effect of First-Dimension Undersampling on Effective Peak Capacity in Comprehensive Two-Dimensional Separations. Anal. Chem. 2008, 80 (2), 461–473. 10.1021/ac071504j

(60) Li, X.; Stoll, D. R.; Carr, P. W. Equation for Peak Capacity Estimation in Two-Dimensional Liquid Chromatography. Anal. Chem. 2009, 81 (2), 845–850. 10.1021/ac801772u

(61) Hagel, L. Peak Capacity of Columns for Size-Exclusion Chromatography. J. Chromatogr. A 1992, 591 (1–2), 47–54. 10.1016/0021-9673(92)80221-F

(62) Basharat, A. R.; Xiong, X.; Xu, T.; Zang, Y.; Sun, L.; Liu, X. TopDIA: A Software Tool for Top-Down Data-Independent Acquisition Proteomics. J. Proteome Res. 2025, 24 (1), 55–64. 10.1021/acs.jproteome.4c00293

(63) LeDuc, R. D.; Taylor, G. K.; Kim, Y.-B.; Januszyk, T. E.; Bynum, L. H.; Sola, J. V.; Garavelli, J. S.; Kelleher, N. L. ProSight PTM: An Integrated Environment for Protein Identification and Characterization by Top-down Mass Spectrometry. Nucleic Acids Research 2004, 32 (Web Server), W340–W345. 10.1093/nar/gkh447

(64) Solntsev, S. K.; Shortreed, M. R.; Frey, B. L.; Smith, L. M. Enhanced Global Post-Translational Modification Discovery with MetaMorpheus. J. Proteome Res. 2018, 17 (5), 1844–1851. 10.1021/acs.jproteome.7b00873

(65) Broadway-Stringer, S.; Jiang, H.; Wadmore, K.; Hooper, C.; Douglas, G.; Steeples, V.; Azad, A. J.; Singer, E.; Reyat, J. S.; Galatik, F.; Ehler, E.; Bennett, P.; Kalisch-Smith, J. I.; Sparrow, D. B.; Davies, B.; Djinovic-Carugo, K.; Gautel, M.; Watkins, H.; Gehmlich, K. Insights into the Role of a Cardiomyopathy-Causing Genetic Variant in ACTN2. Cells 2023, 12 (5), 721. 10.3390/cells12050721

(66) Satomi, Y.; Shimonishi, Y.; Takao, T. N-glycosylation at Asn^491^ in the Asn-Xaa-Cys Motif of Human Transferrin. FEBS Letters 2004, 576 (1–2), 51–56. 10.1016/j.febslet.2004.08.061

(67) Durbin, K. R.; Robey, M. T.; Greer, J. B.; Fellers, R. T.; Bailey, A. O. Fast and Accurate Charge State Deconvolution of Protein Mass Spectra. Anal. Chem. 2025, 97 (28), 14964–14973. 10.1021/acs.analchem.5c00288

(68) Sun, C.; Zhang, W.; Zhou, M.; Myu, M.; Xu, W. Full Window Data-Independent Acquisition Method for Deeper Top-Down Proteomics. Anal. Chem. 2025, 97 (12), 6620–6628. 10.1021/acs.analchem.4c06471

