## Supplementary Information for "An Integrated Experimental and Informatics Strategy Enables Large Proteoform Identification in Top-Down Proteomics"

**Table of Contents**

Supplementary Tables

Table S1. Collisional energy table used for autoMS/MS ……..…………………………3

Table S2. Mass shifts considered for candidate selection…………………………...….3

Table S3. Values used to calculate 2D-LC peak capacity ……...……………..………4

Supplementary Figures

Figure S1. Valve configuration for online SEC-RPLC-MS/MS ……...………………...5

Figure S2. Representative 2D-LC-MS data from the HEPES extraction……………...6

Figure S3. Improving the fidelity of the precursor correction algorithm……………..…7

Figure S4. Representative large proteins identified by TopPIC following precursor mass correction……………………………..…………….………….…………….8

Figure S5. Representative DIA MS/MS spectra for a 63 kDa protein………………..…...9

Figure S6. Comparison of DDA and DIA MS/MS spectra ……………………………...10

Figure S7. Representative large proteins identified using the candidate discovery algorithm and MS/MS validation………………………………………………………11

Figure S8. Identification of NADH-ubiquinone oxidoreductase 75 kDa subunit, mitochondrial (NDUS1, 77 kDa)…………………………………………………………12

Figure S9. Identification of ATP synthase F(1) complex subunit alpha, mitochondrial (ATPA, 55 kDa)………………..……………………………………………………...……13

Figure S10. Identification of pyruvate kinase M1 (PKM1, 58 kDa) ..................…..….14

### Supplementary Tables

**Table S1. Collisional energy table used for autoMS/MS.** A fallback charge state of 15 was used.

| *m/z* | Width | Charge State | Collision Energy (eV) |
|---|---|---|---|
| 500 | 10 | 1 | 35 |
| 500 | 10 | 8 | 20 |
| 500 | 10 | 15 | 18 |
| 1000 | 10 | 1 | 24 |
| 1000 | 10 | 8 | 22 |
| 1000 | 10 | 15 | 20 |
| 1300 | 10 | 1 | 42 |
| 1300 | 10 | 8 | 40 |
| 1300 | 10 | 15 | 38 |
| 2000 | 10 | 1 | 50 |
| 2000 | 10 | 8 | 48 |
| 2000 | 10 | 15 | 45 |

**Table S2. Mass shifts considered for candidate selection.** These mass shifts are used to find candidates in cases where the UniProt annotation may not match the observed proteoform, such as unannotated Met cleavage or N-terminal acetylation.

| Description | Average Mass |
|---|---|
| add Met | 131.1960 |
| remove Met | -131.1960 |
| add acetylation | 42.0368 |
| remove acetylation | -42.0368 |
| add Met, add acetyl | 173.2328 |
| remove Met, remove acetyl | -173.2328 |
| add met, remove acetyl | 89.1592 |

**Table S3. Values used to calculate 2D-LC peak capacity.**

| Extract / Fraction | *m/z* | Apex RT (min) | FWHM (min) |
|---|---|---|---|
| HEPES 12-13 min | 1018.3900; 1031.1400; 1044.1200; 1057.5000; 1071.2000; 1085.3700; 1099.7600; 1114.6000 | 36.57 | 0.30 |
| HEPES 12-13 min | 1004.5186; 1083.2076; 1104.7856; 1150.8182; 1200.9333; 1175.3260 | 42.81 | 0.28 |
| HEPES 12-13 min | 872.6894, 891.2559, 930.778, 951.9518, 997.1588 | 45.42 | 0.33 |
| HEPES 12-13 min | 813.0022, 853.5523, 898.5295, 948.3356, 1004.0593 | 34.76 | 0.20 |
| HEPES 13-14 min | 1034.7500; 1051.9000; 1069.7000; 1088.2000; 1107.3000; 1126.9900; 1147.5000; 1168.7000 | 38.53 | 0.28 |
| HEPES 13-14 min | 1073.0900; 1028.0300; 1074.6800; 1091.2300; 1125.8700; 1143.9700; 1162.7000; 1182.1000 | 40.08 | 0.24 |
| HEPES 13-14 min | 1077.9000; 1094.9000; 1112.3200; 1130.1900; 1148.5000; 1167.7000; 1187.6800; 1207.9000 | 43.03 | 0.23 |
| HEPES 13-14 min | 967.2000; 983.6000; 1000.5000; 1018.0600; 1036.1700; 1055.0300; 1074.5900; 1094.7900 | 44.24 | 0.38 |
| HEPES 14-15 min | 818.7900; 833.3900; 848.4900; 864.2100; 880.4700; 897.4400; 914.9800; 933.2400; 952.3000 | 40.74 | 0.31 |
| HEPES 14-15 min | 1208.9526, 1231.362, 1254.5423, 1278.6444, 1303.7163, 1356.8612 | 37.11 | 0.41 |
| HEPES 14-15 min | 809.0644; 842.6908; 879.2856; 919.2070; 777.9071 | 35.83 | 0.17 |
| HEPES 14-15 min | 857.0619; 885.5961; 916.0992; 948.7820; 983.8109 | 39.86 | 0.23 |
| HEPES 15-16 min | 808.4598; 839.5544; 873.0963; 909.3909; 948.9295 | 43.07 | 0.22 |
| HEPES 15-16 min | 831.6300; 850.9400; 871.1900; 892.4500; 914.7100; 938.1900; 962.7700 | 49.18 | 0.48 |
| HEPES 15-16 min | 932.9990; 991.2489; 1057.2648; 1132.7117 | 32.18 | 0.18 |
| HEPES 15-16 min | 755.9751; 793.7237; 835.4451; 881.8030; 933.5547 | 34.73 | 0.49 |
| HEPES 16-17 min | 739.4400; 778.2500; 821.4300; 869.7500; 923.9900; 985.5200 | 38.53 | 0.29 |
| HEPES 16-17 min | 809.4245; 871.6870; 944.2433; 1258.5429; 755.5972 | 39.87 | 0.20 |
| HEPES 16-17 min | 1123.0791; 1263.3383; 1443.6692; 1010.8722 | 31.8 | 0.17 |
| HEPES 16-17 min | 1262.2948; 1122.0408; 1009.9377; 918.3081 | 45.79 | 0.25 |
| HEPES 17-18 min | 761.9110; 914.0909; 1142.3595 | 37.8 | 0.20 |
| HEPES 17-18 min | 1088.9800; 1224.9800; 1399.9700; 1632.9700 | 47.79 | 0.23 |
| HEPES 17-18 min | 1048.0129, 1197.7276, 1397.1813 | 42.06 | 0.20 |
| HEPES 17-18 min | 721.2870; 757.3996; 797.0519; 841.3328 | 43.48 | 0.21 |
| 1D SEC | 734.1044, 779.7972, 831.6514, 891.0537 | 14.33 | 0.69 |
| 1D SEC | 707.0184; 807.8766; 628.5726 | 14.54 | 0.63 |
| 1D SEC | 1327.4234; 1517.0548 | 14.92 | 0.79 |
| 1D SEC | 1295.5365 | 14.40 | 0.55 |

### Supplementary Figures

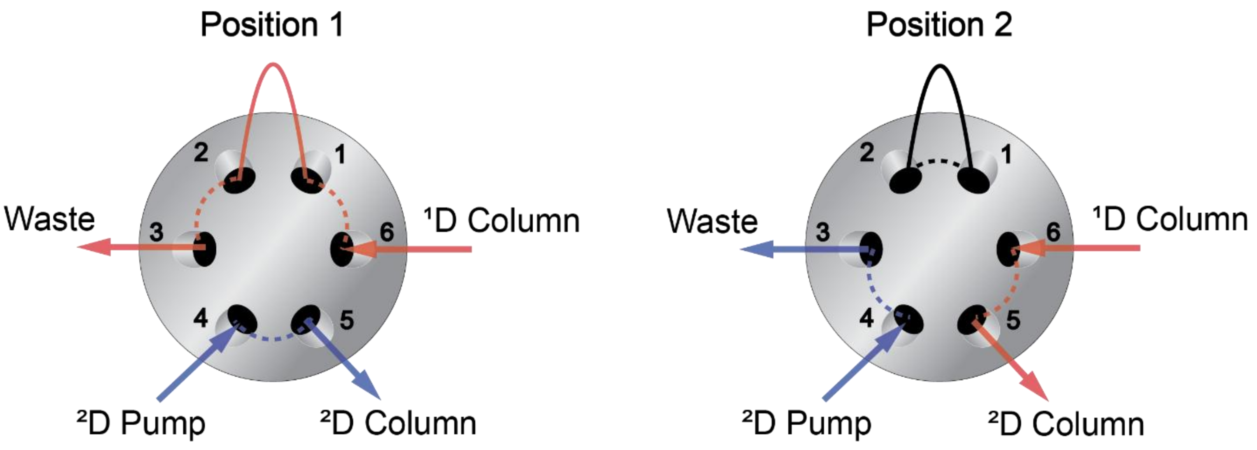


**Figure S1. Valve configuration for online SEC-RPLC-MS/MS.** In position 1, the $^1$D column flows to waste while the $^2$D column can be equilibrated or subjected to a separation gradient. In position 2, $^1$D effluent flows directly into the $^2$D column, trapping proteins eluting from the $^1$D column.

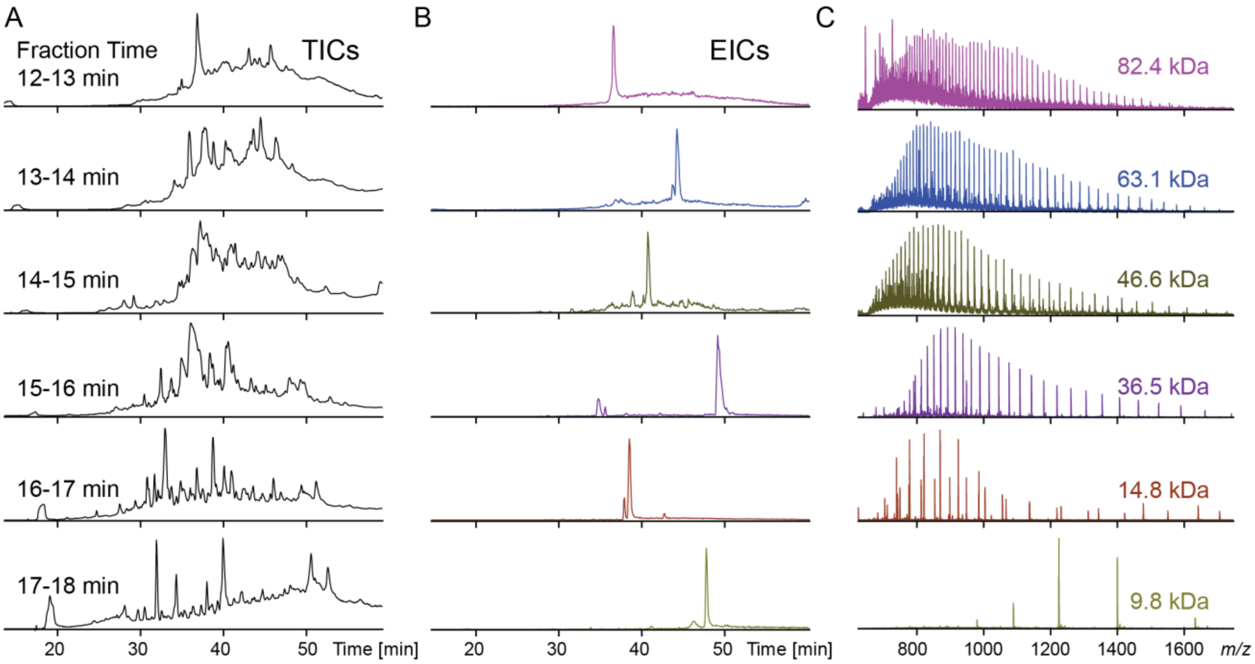


**Figure S2. Representative 2D-LC-MS data from the HEPES extraction. A)** Total ion chromatograms (TICs) for 6 sequential 1-min SEC fractions from a HEPES protein extract. Unique TIC profiles indicate good size-based separation in the first dimension. **B)** Extracted ion chromatograms (EICs) and **C)** mass spectra of representative proteins demonstrating the range of size separation.

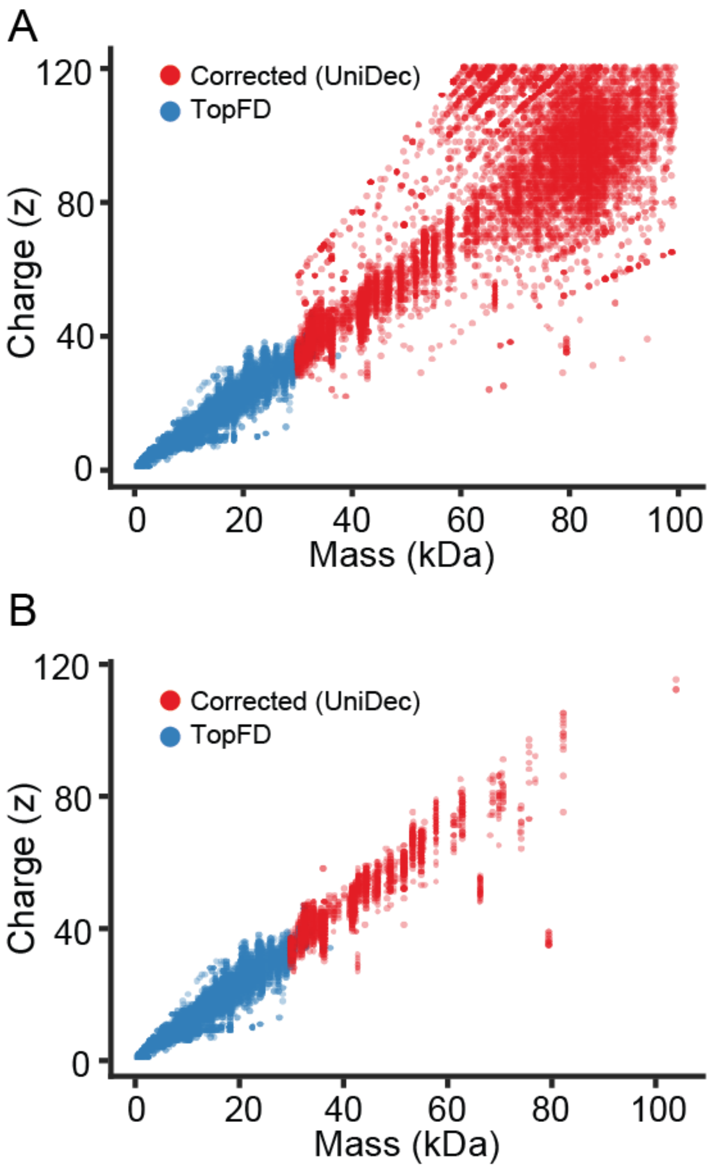


**Figure S3. Improving the fidelity of the precursor correction algorithm. A)** Plot of precursor masses and charge states from initial precursor correction algorithm. **B)** Plot of precursor masses and charge states after adding constraints including a minimum number of isolations and a maximum number of UniDec picked peaks per scan.

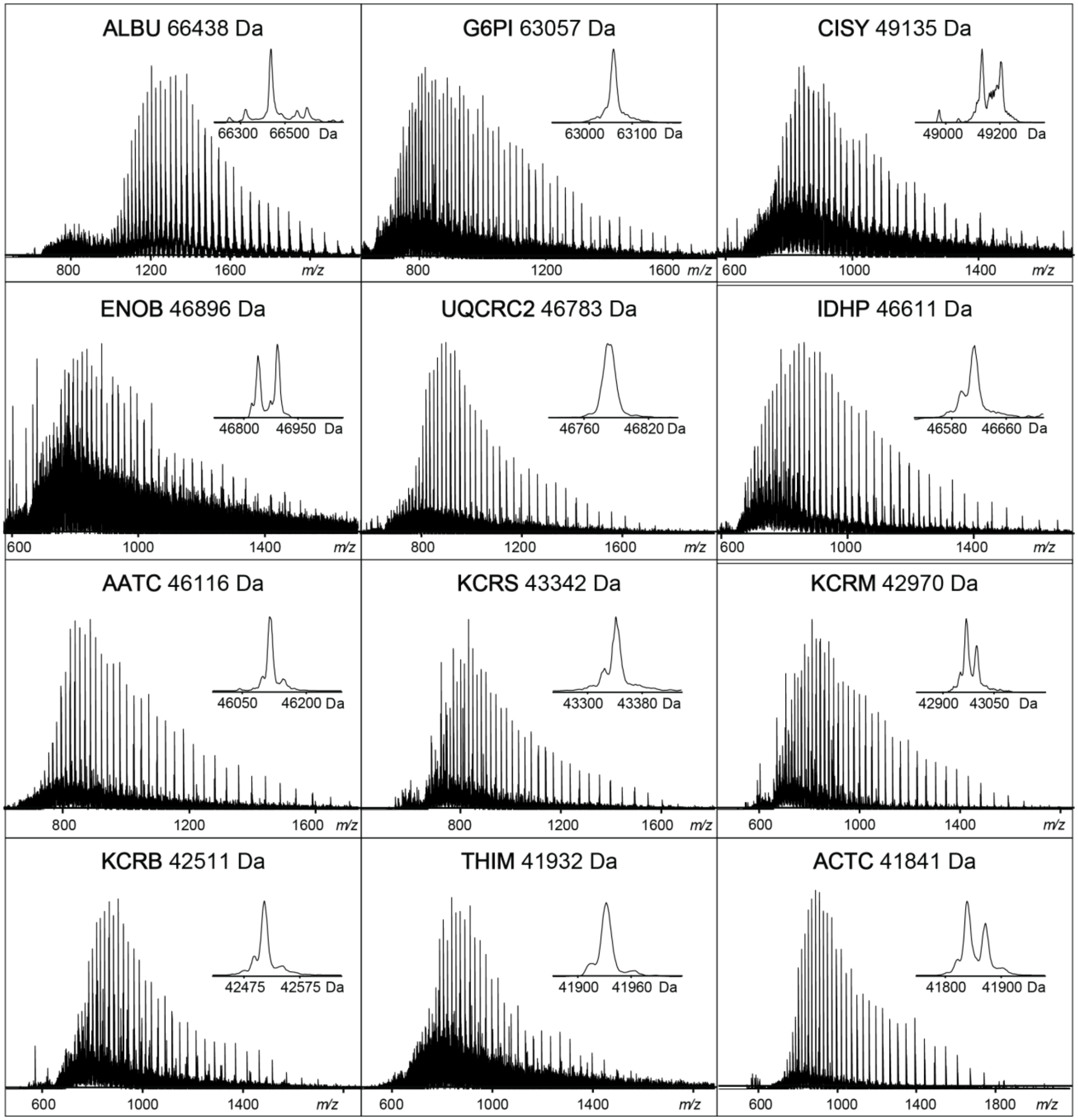


**Figure S4. Representative large proteins identified by TopPIC following precursor mass correction.** Protein symbols correspond to the following proteins: albumin (ALBU), glucose-6-phosphate isomerase (G6PI), mitochondrial citrate synthase (CISY), beta-enolase (ENOB), mitochondrial cytochrome b-c1 complex subunit 2 (UQCRC2), mitochondrial isocitrate dehydrogenase (IDHP), cytoplasmic aspartate aminotransferase (AATC), mitochondrial creatine kinase S-type (KCRS), creatine kinase M-type (KCRM), creatine kinase B-type (KCRB), mitochondrial 3-ketoacyl-CoA thiolase (THIM), and cardiac alpha actin 1 (ACTC).

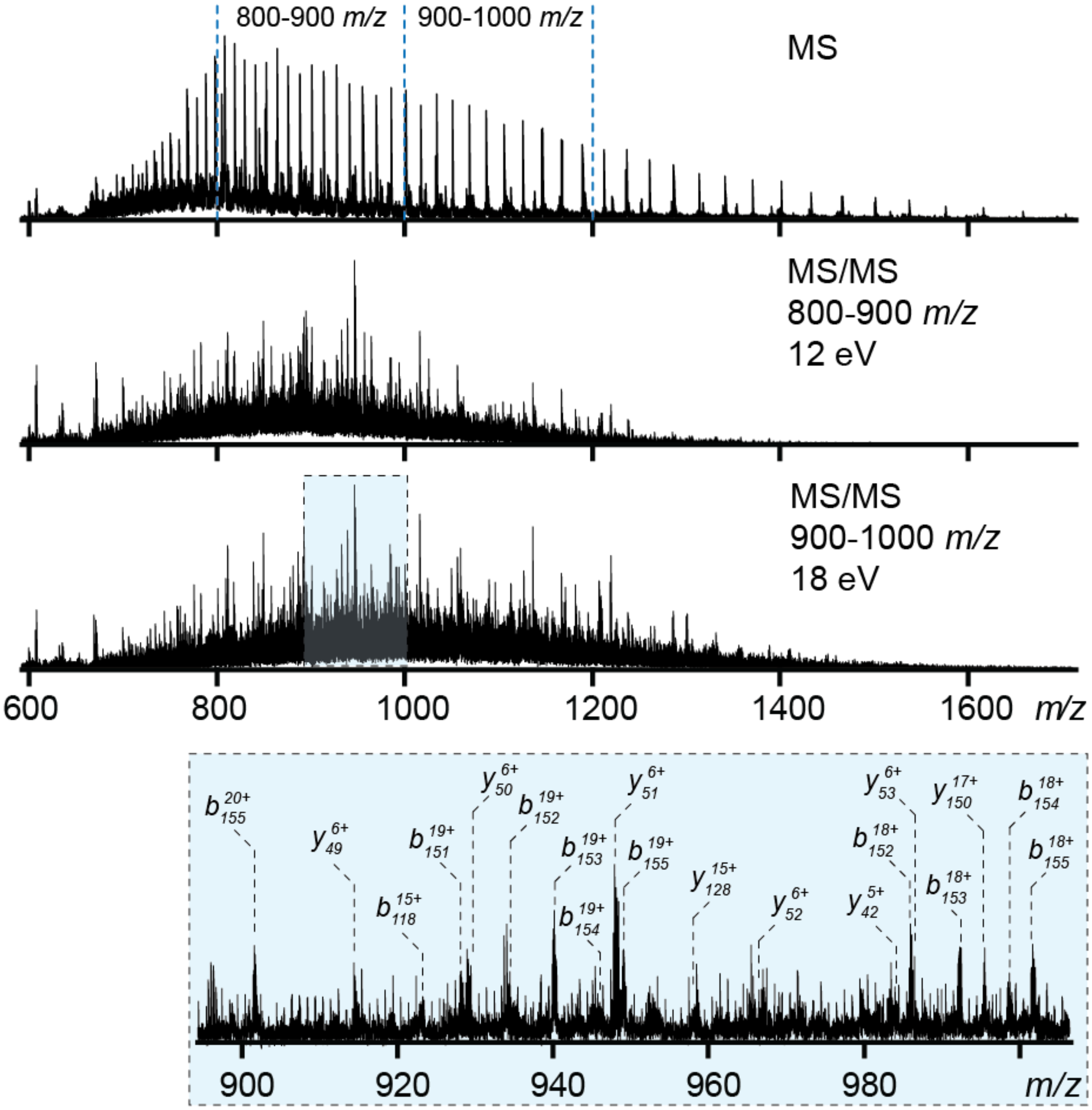


**Figure S5. Representative DIA MS/MS spectral acquisition for a 63 kDa protein.** The DIA method consists of two 100 *m/z* windows to sample the regions containing the most abundant charge states for large proteoforms. This representative example contains data from DIA fragmentation during the elution of glucose-6-phosphate isomerase, a 63 kDa protein, initially identified by TopPIC following precursor correction.

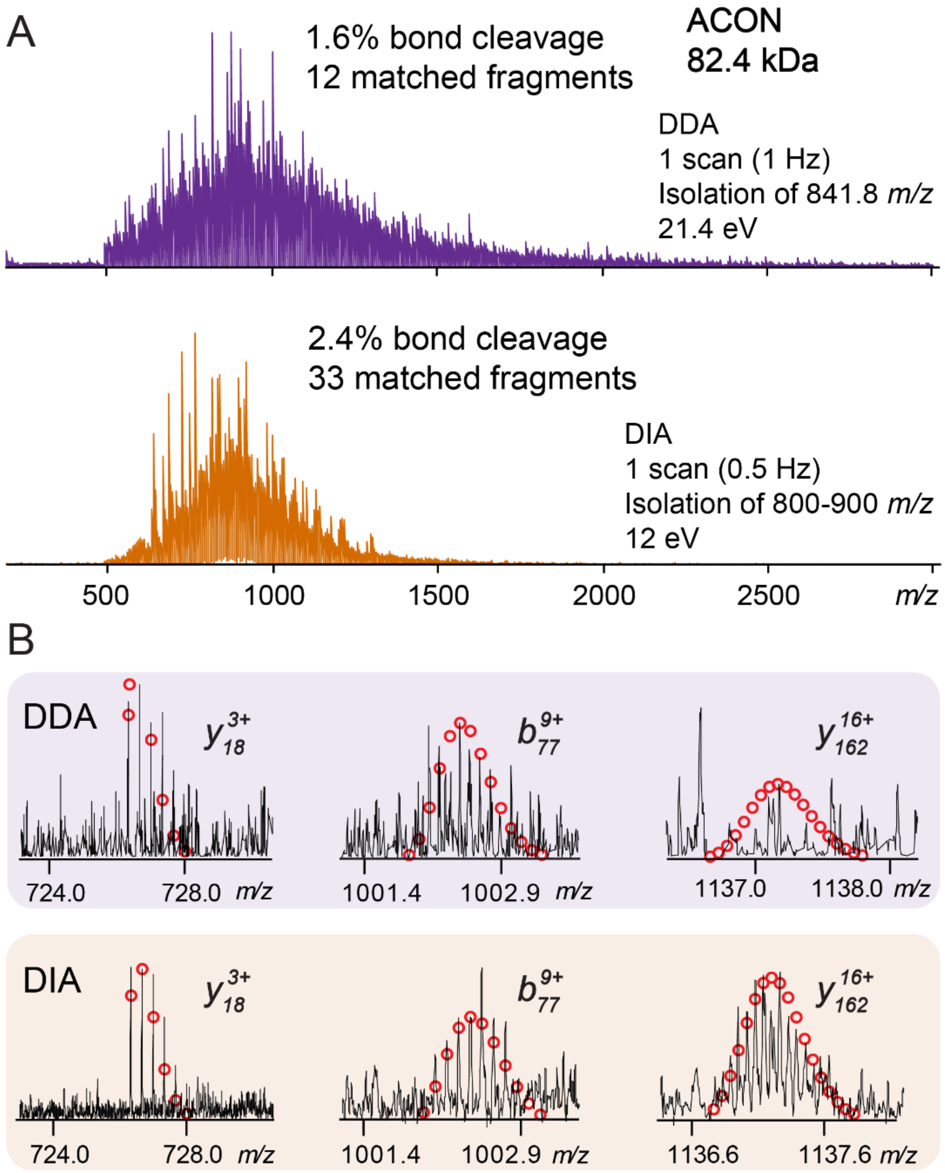


**Figure S6. Comparison of DDA and DIA MS/MS spectra. A)** Example single-scan MS/MS spectra from fragmentation of aconitate hydratase (ACON, 82.4 kDa) by the DDA and DIA method. The DIA method tailored to large proteins achieved higher percentage bond cleavage and results in more matched fragments. **B)** Comparison of fragment ions between the two methods. The DIA method achieves higher signal to noise and detects some fragments which are unobserved in the DDA method.

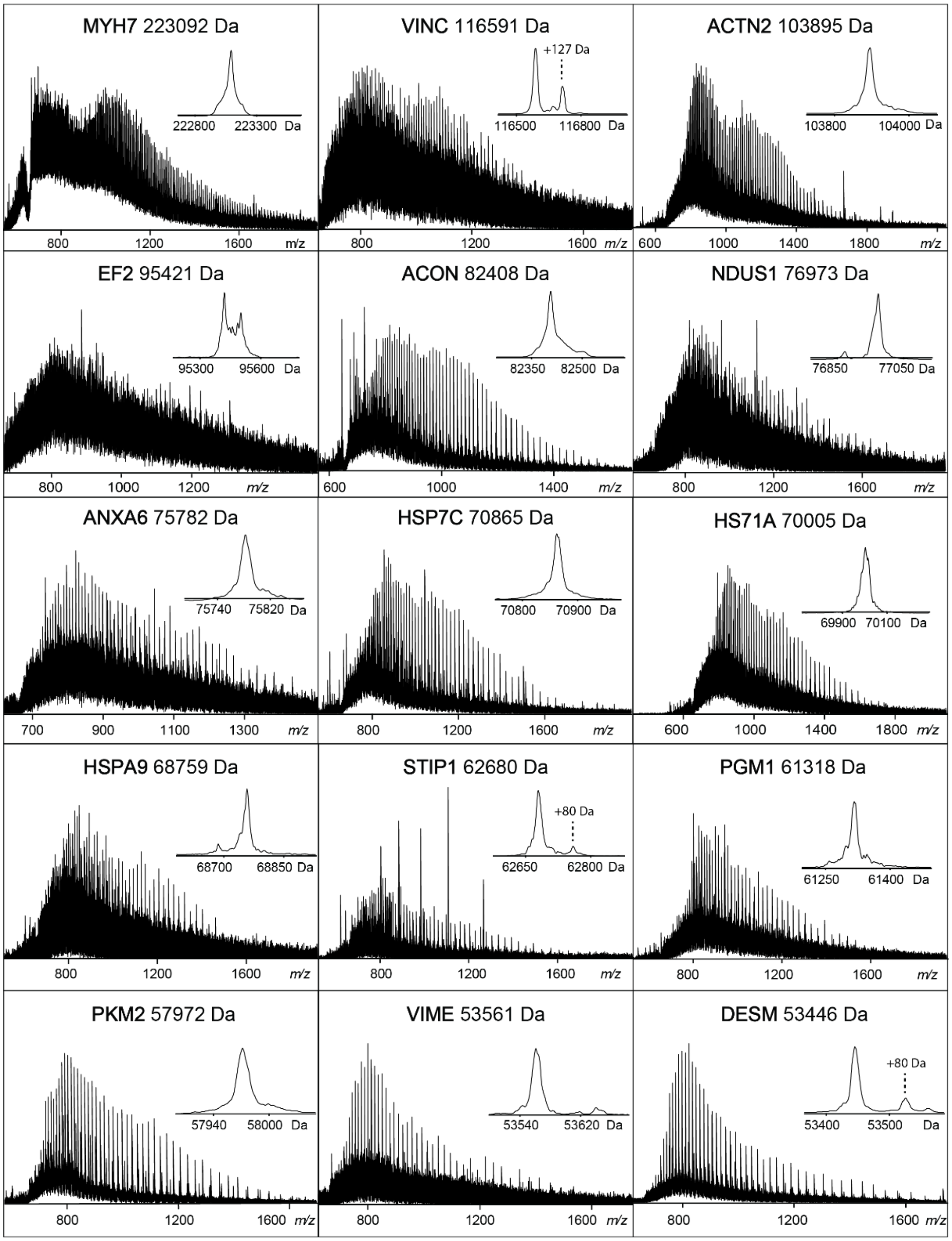


**Figure S7. Representative large proteins identified using the candidate discovery algorithm and MS/MS validation.** Protein symbols correspond to the following proteins: myosin-7 (MYH7), vinculin (VINC), alpha-actinin-2 (ACTN2), elongation factor 2 (EF2), mitochondrial aconitate hydratase (ACON), mitochondrial NADH-ubiquinone oxidoreductase 75 kDa subunit (NDUS1), annexin A6 (ANXA6), heat shock cognate 71 kDa protein (HSP7C), heat shock 70 kDa protein (HS71A), mitochondrial stress-70 protein (HSPA9), stress-induced-phosphoprotein 1 (STIP1), phosphoglucomutase-1 (PGM1), pyruvate kinase M1 (PKM1), vimentin (VIME), and desmin (DESM).

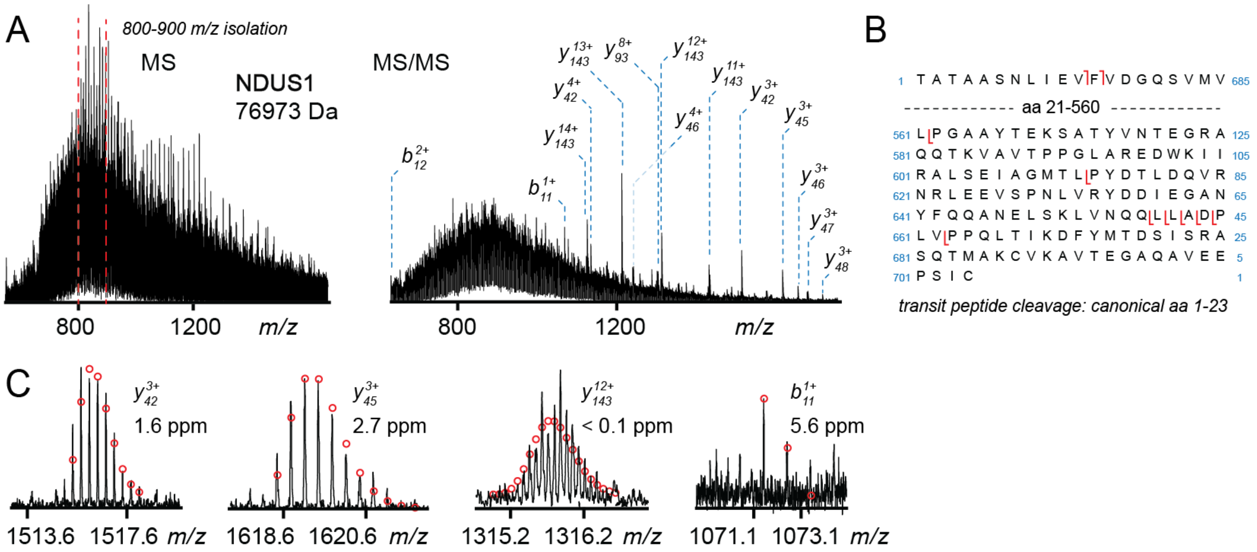


**Figure S8. Identification of NADH-ubiquinone oxidoreductase 75 kDa subunit, mitochondrial (NDUS1, 77 kDa). A)** MS spectrum including annotated DIA isolation window (*red*), and annotated MS/MS spectrum. **B)** Sequence table demonstrating CID fragmentation coverage. **C)** Representative fragment ions with theoretical isotopic fits and high mass accuracy.

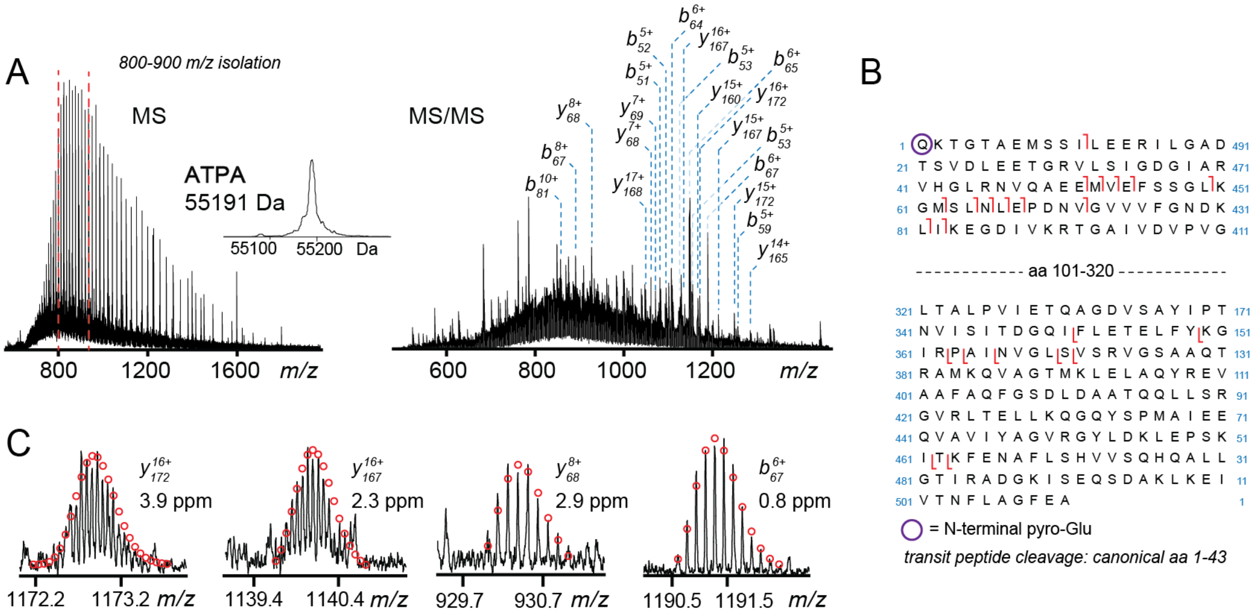


**Figure S9. Identification of ATP synthase F(1) complex subunit alpha, mitochondrial (ATPA, 55 kDa). A)** MS spectrum including annotated DIA isolation window (*red*), and annotated MS/MS spectrum. **B)** Sequence table demonstrating CID fragmentation coverage and discovery of an N-terminal pyro-Glu. **C)** Representative fragment ions with theoretical isotopic fits and high mass accuracy.

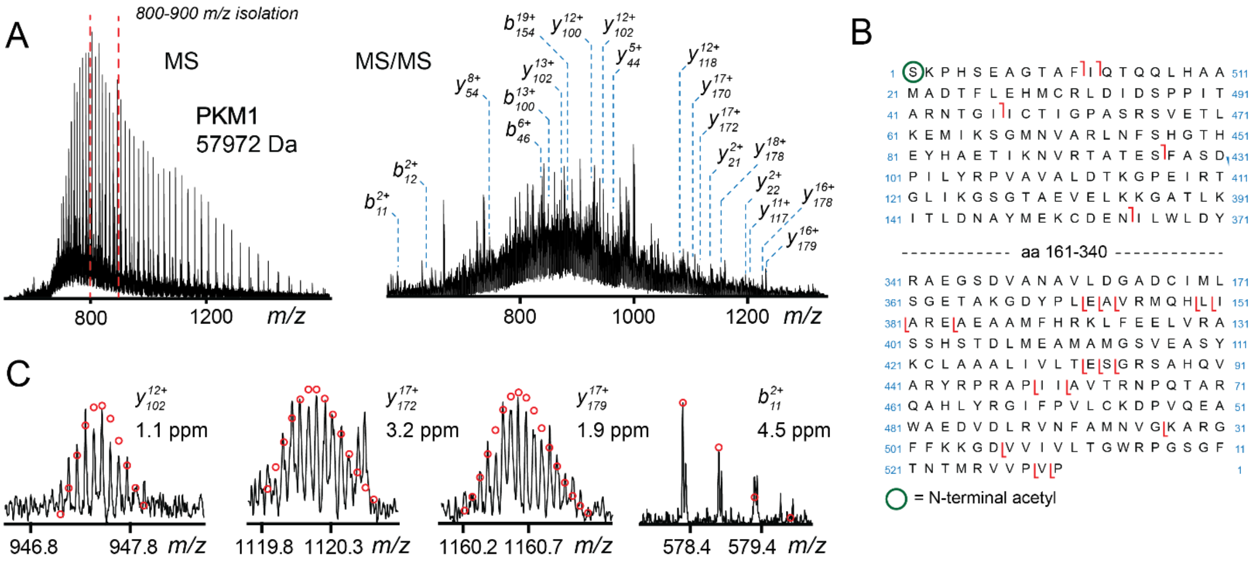


**Figure S10. Identification of pyruvate kinase M1 (PKM1, 58 kDa). A)** MS spectrum including annotated DIA isolation window (*red*), and annotated MS/MS spectrum. **B)** Sequence table demonstrating CID fragmentation coverage. **C)** Representative fragment ions with theoretical isotopic fits and high mass accuracy.
